# Microbial remineralization processes during post-spring-bloom excess phosphate in the northern Baltic Sea

**DOI:** 10.1101/2024.02.02.577174

**Authors:** Mari Vanharanta, Mariano Santoro, Cristian Villena-Alemany, Jonna Piiparinen, Kasia Piwosz, Hans-Peter Grossart, Matthias Labrenz, Kristian Spilling

## Abstract

In the northern Baltic, post-spring-bloom low dissolved inorganic nitrogen to phosphorus conditions, degradation of N-rich organic matter potentially supports the drawdown of excess phosphate. During a 17-day-long mesocosm experiment in the south-west Finnish archipelago, we examined nitrogen, phosphorus and carbon acquiring extracellular enzyme activities in three size fractions (<0.2 µm, 0.2–3 µm, and >3 µm), bacterial abundance, production, community composition and its predicted metabolic functions. The mesocosms received different carbon and nitrogen amendments to test for the effect of inorganic nutrient stoichiometry on enzymatic degradation processes that ultimately determine the export potential of organic matter. Alkaline phosphatase activity occurred mainly in the dissolved form and likely contributed to the excess phosphate conditions. In the beginning of the experiment, peptidolytic and glycolytic enzymes were predicted to be produced by free-living bacteria identified within the classes Actinobacteria and Alphaproteobacteria, whereas the contribution of picocyanobacteria increased towards the end. Our results imply that heterotrophic bacteria lost the competition to picocyanobacteria due to the lack of suitable energy sources. The high hydrolytic rates in fractions <0.2 µm and 0.2–3 µm, found in this study, could potentially retain inorganic nutrients in the surface layer and suppress downward fluxes of organic matter and hence carbon sequestration.

## Introduction

The diatom and/or dinoflagellate dominated spring bloom in the Baltic Sea ends upon the depletion of dissolved inorganic nitrogen (DIN) and, in large parts of the Baltic, typically an excess phosphate concentration of >0.2 μmol L^-1^ remains in the upper water layer (Spilling et al. 2018). This condition is regarded to induce recurring summer blooms of N_2_-fixing cyanobacteria (Niemi 1979, Rahm et al. 2000, Lilover and Stips 2008, Vahtera et al. 2007). However, the main period of cyanobacterial blooms usually starts 2–3 months after the termination of the spring bloom, when the excess phosphate has already been consumed by other organisms (Raateoja et al. 2011, Lips and Lips 2017, Vanharanta and Spilling 2023).

The typical post-spring-bloom low inorganic N:P ratio (DIN:DIP ∼2) indicates strong N-limitation of planktonic productivity (Lignell et al. 2008). This condition likely enhances remineralization of organic nitrogen-rich compounds (Eilola and Stigebrandt 1999) that could allow for the utilization of the excess phosphate pool. The low DIN:DIP ratio may favor the growth of heterotrophic bacteria that have generally low DIN:DIP uptake ratio and high affinity to phosphate due to a relatively low C:P of 45 in their biomass (Kirchman 1994, Nausch et al. 2018). Effective uptake of DIP together with organic forms of nitrogen has the potential to turn the system into combined N and P deficiency (Lignell et al. 2008), but heterotrophic bacteria might be limited by labile carbon-rich substrates (Kirchman 1994, Lignell et al. 2008).

Biological remineralization processes are driven by microbially synthetized extracellular enzymes which hydrolyze high molecular weight organic matter (OM) into smaller substances, thus enabling the acquisition of organic monomers and mineral nutrients into microbial cells. These enzymes exist either attached to their producing cell (i.e., cell-attached) in which case the hydrolysis products are near the cell surface for transport and subsequent intracellular metabolism, or secreted into the water (i.e., dissolved) when hydrolysis products are available also for the non-enzyme producing organisms. Depending on the circumstances, either of these forms of OM hydrolysis can be optimal (Traving et al. 2015), and different phyla of heterotrophic bacteria possess varying strategies of enzyme production (Allison et al. 2014, Murray et al. 2007, Ramin and Allison 2019). Different forms of substrate hydrolysis can also depend on the phytoplankton bloom type (Grossart et al. 2007) or phase and environmental conditions, and this can be either due to community change or ability of individual cells to shift between substrate hydrolysis strategies (Traving et al. 2022) that could be related to their ability to alternate between free-living and surface-attached stages (Grossart 2010).

Dissolved enzymes can originate from direct release by prokaryotes attached to particles or colloids in low diffusion and high substrate environments (Baltar et al. 2010, Grossart et al. 2007, Ziervogel and Arnosti 2008, Ziervogel et al. 2010), bacterial stress and mortality (Baltar et al. 2019), cell lysis due to viral infection (Karner and Rassoulzadegan 1995) and protist grazing on bacteria (Bochdansky et al. 1995). Also, fungi (Grossart et al. 2019, Salazar-Alekseyeva et al. 2023) and autotrophs such as diatoms (e.g. Rengefors et al. 2001) and mixotrophic algae (Salerno and Stoecker 2009) produce extracellular enzymes that, e.g. during phytoplankton bloom breakdown can end up in the dissolved enzyme pool. In the Baltic Sea, alkaline phosphatase (APase) activity has been attributed to phytoplankton (Nausch 1998) and is often found in the cell-attached form (González-Gil et al. 1998, Strojsová et al. 2003), whereas up to 30 % of the bacterial APase activity appears to be dissolved (Luo et al. 2009). In addition, physical variables such as solar irradiation (Thomson et al. 2017) and low temperature (Steen and Arnosti 2011, Baltar et al. 2016) have been associated with a high contribution of dissolved extracellular enzyme activity (EEA) to total hydrolytic activity. The ecologically significant characteristic of dissolved enzymes is that they may remain functional in the water column for days to weeks after being released into the surrounding water (Hoppe 1991, Ziervogel et al. 2010, Steen and Arnosti 2011, Thomson et al. 2019). Thereby, they spatially and temporally decouple hydrolysis products and enzyme producing cell (Ziervogel et al. 2010).

In a recent indoor tank experiment, a larger part of the excess phosphate remaining after the spring bloom ended up in the particulate organic phosphorus pool than in the dissolved organic fraction (Vanharanta and Spilling 2023). While the dissolved organic matter pool would increase internal cycling in the upper water layer, the particulate form might be heavily colonized by bacteria (Bochdansky et al. 2016) and has a higher potential to sink out from the surface layer, thereby transporting the EEA attached to it to the seafloor. In contrast, EEA present in the dissolved form or attached to free-living cells would more likely retain nutrients and other hydrolysis products suspended in the surface layer or cycle in the microbial food web (Hoppe et al. 2002), respectively. Studying EEAs in size fractionated samples can therefore give insights to particle export and transfer efficiency (Karrach et al. 2003).

As part of a larger study on the fate of excess phosphate in the northern Baltic Sea, the present paper reports the enzymatic processing of OM under low DIN:DIP ratio relative to the canonical N:P Redfield ratio 16:1 (Redfield 1958). Inorganic nutrient supply and availability of labile carbon sources influence the uptake stoichiometry and resource competition between primary producers and heterotrophic bacteria with consequences for further trophic interactions (Thingstad et al. 2008). The main aim of this study was to investigate the possible drivers of EEAs and the degradation potential of organic material that ultimately determines the potential of OM export to the sea floor after the northern Baltic spring bloom. In addition, we described the bacterial community composition and compared the measured EEAs with the predicted functional abundances of the bacterial taxa identified via 16S rRNA gene sequences. We hypothesized that (1) DIN limitation and excess phosphate availability enhance the degradation of dissolved organic nitrogen (DON) enabling the removal of the excess phosphate pool and consequently could drive the system into co-deficiency of DIN and DIP; (2) heterotrophic bacteria may be also limited by labile carbon substrates which reduces their competitive edge against picophytoplankton for inorganic nutrients; and (3) high rates of enzyme activity contribute to OM dissolution reducing the potential for downward fluxes of particulate OM (POM).

## Materials and methods

### Experimental design and sampling

An outdoor mesocosm experiment was conducted between 7^th^ and 23^rd^ June 2021. Scientific foundation and details of the experimental setup are given in Spilling et al. (REF). Briefly, we moored 14 transparent plastic mesocosm bags (volume 1.2 m^3^, diameter 0.9 m, depth 2 m) in a linear array beside a pontoon off Tvärminne Zoological Station at the Finnish southwest coast (59° 50’ 37” N, 23° 15’ 6” E). Mesocosms were filled on 7^th^ June marking day -1 of the experiment. Temperature and salinity were ∼18℃ and 5.5 psu, respectively. Dummy bags filled at each end ensured the same light conditions in mesocosm 1 and 12. All mesocosms were partially protected from rain and bird feces with a transparent plastic roof.

On experiment day 0 (8^th^ June), the concentration of DIP was 0.013 µmol L^-1^ and each of the 12 mesocosms received an addition of PO_4_^3-^ (KH_2_PO_4_) to a starting concentration of 0.66 µmol L^-1^ to simulate the excessive DIP conditions prevailing in the Gulf of Finland after the spring bloom. In addition to three otherwise unamended control mesocosms, the experimental design consisted of three different triplicate treatments: carbon addition as glucose (C-treatment), nitrate addition (N-treatment), and combined carbon and nitrate additions (NC-treatment). Before the start of the experiment there was 0.031 µmol DIN L^-1^ in the water. Nitrate (NO_3_^-^) was added in each of the three N- and NC-treated mesocosms to a final starting concentration of 3.66 µmol NO_3_^-^ L^-1^. Glucose was added to each of the three C- and NC-treated mesocosms to final concentration of 36 µmol C L^-1^. All additions were made on day 0, right before the first full sampling.

Water samples for extracellular enzyme activities, chlorophyll-*a*, inorganic nutrient, particulate and dissolved organic nutrient concentrations, bacterial abundance, and production, as well as DNA extraction for downstream 16S rRNA analysis were taken on days 0, 1, 3, 6, 8, 10, 13, and 15 with a Limnos sampler (Hydro-Bios, Germany) at 1.5 m depth. Temperature was measured inside the mesocosms by logging sensors (HOBO U26; Onset Inc, US) throughout the entire experiment. Water collected from each mesocosm was transported in closed plastic containers, previously pre-rinsed with the sampled water, to a climate chamber in the laboratory for further processing.

### Chemical analyses

Dissolved inorganic nutrients (NO_3_^-^+NO_2_^-^, PO_4_^3-^) were measured by standard colorimetric methods (Grasshoff et al. 1999) using a photometric analyzer (Thermo Scientific Aquacem 250). NH_4_^+^ was measured with a Hitachi U-1100 spectrophotometer.

Chlorophyll-*a* concentration was determined according to Jespersen and Christoffersen (1987) by filtering 50 mL of sample in duplicates onto GF/F filters (pore size 0.7 µm, Whatman) and extracted with 94 % ethanol in the dark at room temperature for 24 h before analysis with a fluorometer (Varian Inc., Cary Eclipse). Chlorophyll*-a* measurements were calibrated with a chlorophyll-*a* standard (Sigma).

Samples for dissolved organic carbon (DOC) and total dissolved nitrogen (TDN) (20 mL) were filtered into acid-washed and pre-combusted glass vials through 0.2 µm polycarbonate syringe filters (Whatman). Samples were acidified with 2 M HCl and concentrations were determined using a Shimadzu TOC-VCPH analyzer equipped with a chemiluminescence detector (Shimadzu TNM-1) for measuring TDN. DON was calculated as the difference between TDN and DIN. Total dissolved phosphorus (TDP) concentration was determined according to Koistinen et al. (2017) from 30 mL of sample water filtered through 0.2 µm polycarbonate syringe filters (Whatman) into acid-washed centrifuge tubes. Dissolved organic phosphorus (DOP) was calculated as the difference between TDP and DIP.

For determination of particulate organic nutrient concentrations, 100 mL of water was filtered in duplicates onto acid washed (2 M HCl) and pre-combusted (450°C, 4 h) GF/F filters (Whatman). Concentrations of particulate organic carbon (POC) and nitrogen (PON) were measured with a CHN element analyzer coupled to a mass spectrometer (Europa Scientific). Concentration of particulate organic phosphorus (POP) was measured by dry incineration and acid hydrolysis (Solórzano and Sharp 1980) with modification by Koistinen et al. (2017). Particulate organic elemental ratios POC:PON:POP were calculated on a molar basis.

### Biovolume estimates

The microphytoplankton composition, total biovolume, and detritus biovolume were determined by FlowCam (see further methodological details in Spilling et al. REF). The contribution of detritus to the total biovolume was used as an indicator of OM quality.

### Bacteria abundance and bacterial production

Heterotrophic bacteria were enumerated by flow cytometry (LSR II, BD Biosciences, USA) using a 488 nm laser. Samples were fixed with 1 % paraformaldehyde (final concentration) for 15 min in darkness, flash frozen in liquid nitrogen, and stored at -80℃ until analysis. Prior to flow cytometry, samples were stained with SYBRGreen I (Molecular Probes, Eugene, OR, USA) at a 10^-4^ (v/v) concentration and incubated for 15 min in the dark according to Gasol and Del Giorgio (2000). CountBright beads (Molecular Probes) were added to each sample to determine measured volume. Heterotrophic bacteria were detected according to their green fluorescence and side scatter properties using the FACSDiva Software (BD Biosciences). Cell counts were obtained with the Flowing Software version number 2.5.1 (https://bioscience.fi/?s=flowing+software).

Bacterial production was measured as incorporation of ^3^H-thymidine (BPT) and ^14^C-leucine (BPL) using cold trichloroacetic (TCA) extraction (Fuhrman and Azam 1982, Kirchman et al. 1985). Of each sample three replicates and two formaldehyde-fixed (final conc. 1.85 %) adsorption blanks were spiked with [methyl-^3^H]-thymidine and L-[^14^C(U)]-leucine (Perkin Elmer) at saturating concentrations of 20 nM and 150 nM, respectively. The sub-samples (1 mL) were incubated in Eppendorf tubes for 1–1.5 h in the dark at ∼17°C and the incubation was stopped by addition of formaldehyde (fin. conc. of 1.85 %). TCA extraction was done using the centrifugation method (Smith and Azam 1992) after which the pellets were dissolved in Instagel scintillation cocktail for measurements with a Wallac Win Spectral 1414 liquid scintillation counter. A cell conversion factor of 1.4 ✕ 10^9^ cells nmol^-1^ (HELCOM 2008) and a carbon conversion factor of 0.12 pg C × (µm^3^ cell^-1^)^0.7^ (Norland 1993) were used to convert thymidine incorporation to carbon production (µg C l^-1^ h^-1^). Based on earlier measurements an average bacterial cell volume of 0.06 µm^3^ was used (Hoikkala pers. comm.). Leucine incorporation was converted to carbon production using a factor of 1.5 kg C mol^-1^ (Simon and Azam 1989).

### Extracellular enzyme activities

Hydrolytic rates of four enzymes; alkaline phosphatase (APase), β-glucosidase (BGase), α-glucosidase (AGase) and leucine aminopeptidase (LAPase) were measured as an increase in fluorescence over a 4 h incubation in the dark at ∼17°C. Fluorescence was measured in 1 h intervals with an Agilent Cary Eclipse Fluorescence Spectrophotometer with excitation/emission wavelengths of 360/450 nm on Nunc 96 well plates. Working solutions of fluorogenic model substrate analogs corresponding to classes of natural substrate chemical bonds, 4-methylumbelliferyl-phosphate (APase), 4-methylumbelliferyl-βD-glucopyranoside (BGase), 4-methylumbelliferyl-αD-glucopyranoside (AGase), and L-leucine-7-amido-4-methylcoumarin (LAPase) (Sigma, Aldrich) were prepared according to Hoppe (1983) and added in samples at previously tested saturating final concentrations of 100 µM (APase, BGase, AGase) and 500 µM (LAPase). Consequently, measured hydrolytic rates represent potential values proportional to the pool of extracellular enzymes present in the sample (Hoppe 1983). Substrate analogs were measured in four replicates per sample. Fluorescence was converted to microbial enzymatic activity (nmol L^-1^ h^-1^) by standard curves established with a range of increasing concentrations of chromophores 4-methylumbelliferone and 7-amido-4-methylcoumarin added in 0.2 µm filtered sample water according to Baltar et al. (2016). Autoclaved water samples as controls have previously indicated negligible background activity.

Sample water for EEA measurements was size fractionated by filtering a subsample through sterile 0.2 µm low protein binding Acrodisc filters (Pall) and another subsample through 3 µm Nuclepore Track-Etch Membrane filters (Whatman). The <0.2 µm fraction is assumed to contain free dissolved enzymes. The EEA in size fraction 0.2–3 µm was obtained from the difference in activities in 0.2 µm and 3 µm filtrates and mainly represents extracellular enzymes attached to free-living bacteria or dissolved enzymes attached to small abiotic surfaces. In addition, activity in unfiltered sample water was measured for obtaining the >3 µm activity by calculating the difference in activities in the non-size fractionated sample water and the <3 µm filtrate. This fraction represents either particle-attached microbes, dissolved enzymes absorbed to abiotic particles or enzymes produced by large phytoplankton cells. Bacteria-specific EEA (amol cell^-1^h^-1^) was calculated for LAPase, AGase and BGase activities by dividing the respective EEA rate in the 0.2–3 µm fraction by the respective bacterial abundance.

### Bacterial community composition and prediction of metabolic functions

Subsamples of 500 mL of water were filtered onto sterile polycarbonate filters in replicates (Whatman® Nuclepore™ Track-Etched Membranes diam. 47 mm, pore size 0.2 μm). Filters were placed in sterile cryogenic vials before immediate flash-freezing in liquid nitrogen and stored at -80℃. DNA extraction followed the protocol from Nercessian et al. (2005) with slight modifications. V3–V4 region of the 16S rRNA gene were amplified using the primer set 341F– 785R (Klindworth et al. 2013) and sequenced using Illumina MiSeq 2 x 300 bp (Illumina Inc., San Diego, CA, United States). Demultiplexed and adapter clipped reads obtained from LGC Genomics were processed as mixed ones. Sequence data processing was performed according to a tailored snakemake-implemented DADA2 workflow with specific parameter settings provided as supplementary material. 16S rRNA gene copy number correction was performed for all sequences with PICRUSt2 (Phylogenetic Investigation of Communities by Reconstruction of Unobserved States 2; v.2-2.5.1). This approach aimed to use 16S sequences information to predict gene families present in the bacterial community and to combine them in order to estimate the composite metagenome in a tailored workflow (Callahan et al. 2016, Douglas et al. 2020, Köster et al. 2012), including the following steps: (i) maximum likelihood-based phylogenetic placement of sequences on a user-supplied reference tree and alignment via EPA-NG v0.3.8 (Evolutionary Placement Algorithm previously implemented in RAxML, Barbera et al. 2019), (ii) hidden-state prediction for 16S rRNA gene copy, Enzyme Commission (EC) numbers, and Kegg Orthologs (KO) abundances per-genomes, (iii) metagenome prediction, predicted marker gene abundances and predicted gene family abundances from the amplicon sequence variants (ASVs) abundance table. The input ASVs abundance table (containing sequence abundances in read counts) was normalized by the predicted number of 16S rRNA gene copies known for each taxon. The predicted functional profiles per sample were then determined. The normalized sequence abundance table and the weighted nearest-sequenced taxon index (NSTI) values per-sample were obtained. All ASVs with NSTI > 1 were excluded from the downstream analysis. ASVs contributions were included in the output file consisting of the table with ASVs normalized by predicted 16S rRNA gene copy number abundances and predicted metabolic functions with enzymes identified by EC numbers. The ASVs occurring with a minimum relative abundance of 5 % over the whole community were subsetted and then their contributions to the predicted alkaline phosphatase (EC:3.1.3.1), α-glucosidase (EC:3.2.1.20), β-glucosidase (EC:3.2.1.21) and leucyl aminopeptidase (EC:3.4.11.1) were investigated.

### Data analysis

Linear mixed effects modeling (LME) was applied to determine effects on enzymatic activities in different size fractions. Fixed explanatory variables included temperature, chlorophyll-*a*, inorganic and organic nutrient concentrations, and the elemental ratios of organic nutrients. In addition, the effect of treatment was investigated separately for each enzyme in each size fraction. For activities of AGase, BGase, and LAPase in the bacterial size fraction (0.2–3 µm), cell-specific activities were used as response variable and cell-specific bacterial production was included as an additional fixed effect. Mesocosm was used as the random effect. Auto-correlation function (ACF) was used to detect patterns of autocorrelation, and an appropriate autocorrelation structure was included in the model when appropriate. Model assumptions were verified by examination of residuals and the response variable was square root or log_10_-transformed prior to model refitting when required to ensure normality and homogeneity. Akaike information criteria (AIC) was used in model selection and optimal models were selected based on restricted maximum likelihood estimation (REML) criterion. Results were considered significant at p <0.05. Mixed effects modeling was conducted in R v.4.3.0 (R Core Team, 2022) using package nlme (Pinheiro et al. 2023) and package emmeans (Length 2023) was used to obtain estimated marginal means for mixed effects models investigating the treatment effect. Graphs were done in the R package ggplot (Wickham 2016) and in base R with occasional post-processing in Inkscape (Inkscape Project, 2020).

## Results

### Temperature, chlorophyll, and inorganic nutrients

The water temperature was relatively high at the start of the experiment (∼18°C). There was a temporary 5.3°C decline between days 5 and 9 caused by upwelling of cold water that cooled water inside the mesocosm bags to about 13°C. Thereafter, the temperature gradually increased to reach >20°C on the last day of the experiment (Fig. 1A). There was a steep decrease in chlorophyll-*a* concentration from ∼3.4 to ∼0.8 µg L^-1^ between days 0 and 3 in the control and C-treatment, and from ∼3.9 to ∼2.1 µg L^-1^ between days 1 and 3 in the N-and NC-treatments (Fig. 1B) which coincided with a depletion of NO_3_^-^ (Fig. 1C). The chlorophyll-*a* concentration increased again between days 6 and 15, especially in the C-treatment. Compared to the C-treatment and the control, phosphate concentration (P) decreased faster in the N- and NC-treatments, but P was not depleted in any treatment during the entire experiment (Fig. 1D). The remaining P at the end of the experiment was ∼0.16 µmol L^-1^ and ∼0.35 µmol L^-1^ in the mesocosms with N amended (N- and NC-treatment) and unamended (control and C-treatment) mesocosms, respectively. The fastest phosphate uptake rate occurred during the first three days of the experiment during which the added nitrate became depleted in all treatments indicating active phytoplankton growth. The ammonium concentrations declined by almost half in all mesocosms after day 1 from ∼0.32 ± 0.02 µmol L^-1^ to ∼0.17 ± 0.05 µmol L^-1^ on day 15 (Fig. 1E).

**Fig 1.**
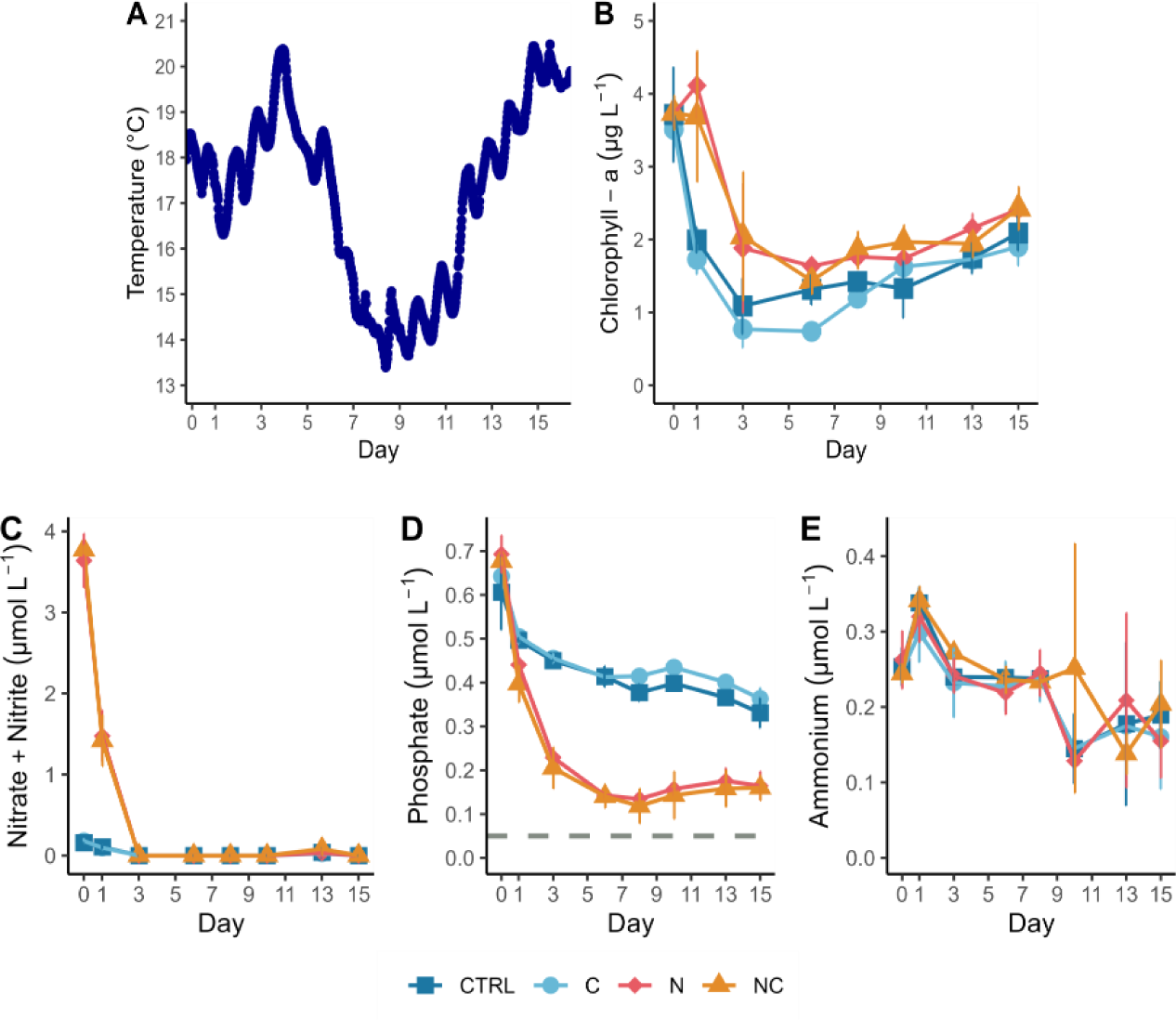
(A) Temperature logger data from mesocosm 1 and concentration time series of (B) chlorophyll-*a*, (C) nitrate+nitrite (µmol NO_3_^-^+NO_2_^-^ L^-1^), (D) phosphate (µmol PO_4_ ^3-^ L^-1^) and (E) ammonium (µmol NH_4_ ^+^ L^-1^). Error bars represent standard deviations of triplicate mesocosm bags. Non-visible error bars are smaller than the marker. Dashed horizontal line in (C) indicates concentration of phosphate depletion (0.05 µmol L^-1^).

### Dissolved and particulate organic nutrients

The dissolved organic nutrient concentrations did not vary among treatments (Fig. 2A–C). DOC concentrations were relatively stable throughout the experiment (∼0.52 ± SD 0.05 mmol L^-1^), whereas DON concentrations showed a pattern of production and removal. DON increased in all treatments during the first three days of the experiment (Fig. 2B), when the uptake of nitrate+nitrite was fast (Fig. 1C). At the end of the experiment, on average, DON concentration was 3.6 µmol L^-1^ higher compared to the start of the experiment, indicating accumulation of refractory DON. The DOP concentration increased from ∼0.36 ± 0.03 µmol L^-1^ to ∼0.45 ± 0.04 µmol L^-1^ between days 1 and 6, after which the DOP decreased close to the starting concentration ∼0.35 ± 0.03 µmol L^-1^ at the end of the experiment (Fig. 2C).

**Fig. 2.**
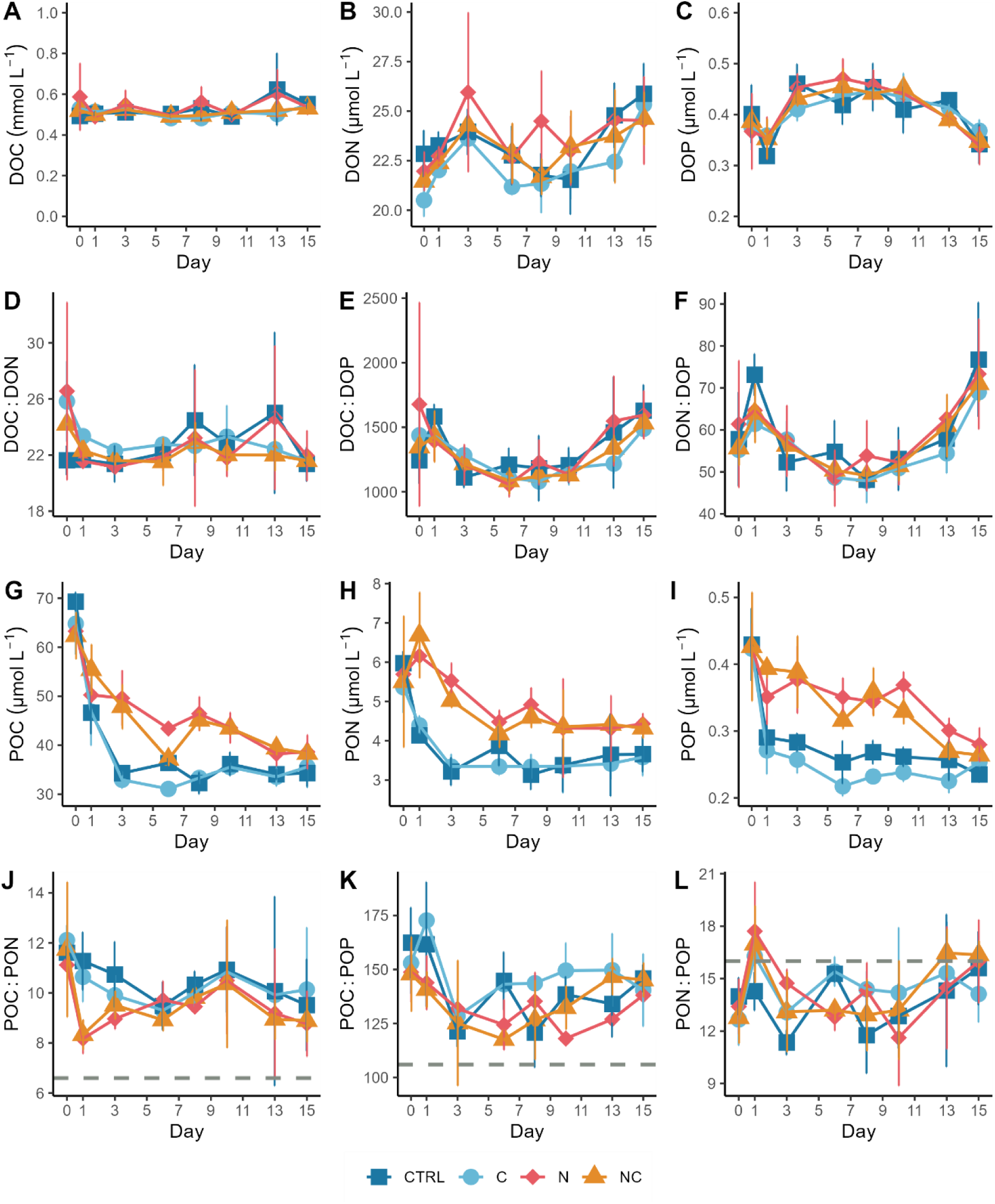
Concentrations of (A) dissolved organic carbon (DOC), (B) dissolved organic nitrogen (DON), (C) dissolved organic phosphorus (DOP), and (D, E, F) stoichiometric ratios of dissolved organic nutrients over time. (G) Particulate organic carbon (POC), (H) particulate organic nitrogen (PON), (I) particulate organic phosphorus (POP) and (J, K, L) stoichiometric ratios of particulate organic nutrients over time. Horizontal gray dashed lines in panels J–L indicate molar Redfield ratios. Error bars represent standard deviations of triplicate mesocosm bags.

The DOC:DON ratio ranged between 17 and 31 without any clear differences among treatments (Fig. 2D). Both DOC:DOP and DON:DOP ratios decreased until the mid-experiment and thereafter increased similarly in all treatments towards the end of the experiment (Fig. 2 E,F).

Particulate organic nutrient concentrations were higher in the N- and NC-treatments compared to the control and C-treatments from day 1 until the end of the experiment (Fig. 2G–I). In the control and C-treatment, the steepest decrease of all particulates occurred during the first half of the experiment after which the concentrations stabilized during the latter phase of the experiment. For both POC and POP in the N- and NC-treatments, the decrease was slower compared to the C-treatment and the control but constant throughout the entire experiment. The PON concentration decreased during the first three days of the experiment as the DON concentration increased, and the uptake of DIN was fast.

POC:PON and POC:POP ratios were higher than the Redfield values 6.6 and 106, respectively, while PON:POP ratios were below the Redfield value (16) almost throughout the entire experiment (Fig. 2J–L).

The contribution of detritus to the total biovolume generally decreased in all treatments during the first half of the experiment from ∼54.1 % to ∼27.7 % after which it increased again towards the end of the experiment (Fig. S1). Lowest mean contribution of detritus to the total biovolume was observed in the NC-treatment on day 8 (18.6 %).

### Bacterial abundance and production

Bacterial growth occurred in all mesocosms (Fig. 3A). Different treatments started to show varying effects on heterotrophic bacteria after day 3 and the abundance peaked on day 6 (C-addition) or day 8 (other treatments), which was 2–5 days after the bacterial production maximum (Fig. 3B,C). The highest average abundances of heterotrophic bacteria occurred in treatments with N- and NC-additions, 7.4×10^6^ and 6.6×10^6^ cells mL^-1^, respectively. A sharp drop in bacterial abundance occurred soon after the peak and the bacterial abundance was generally similar during the first and last samplings. Bacterial production increased in all treatments after an initial drop between the start and day 1 but was lower in the control compared to all other treatments (Fig. 3B,C). After the peak on day 6, bacterial production decreased to lower levels than at the start of the experiment. BPL stagnated at slightly higher level in N and NC-treatments whereas the treatment effect in BPT had faded by day 13.

**Fig. 3.**
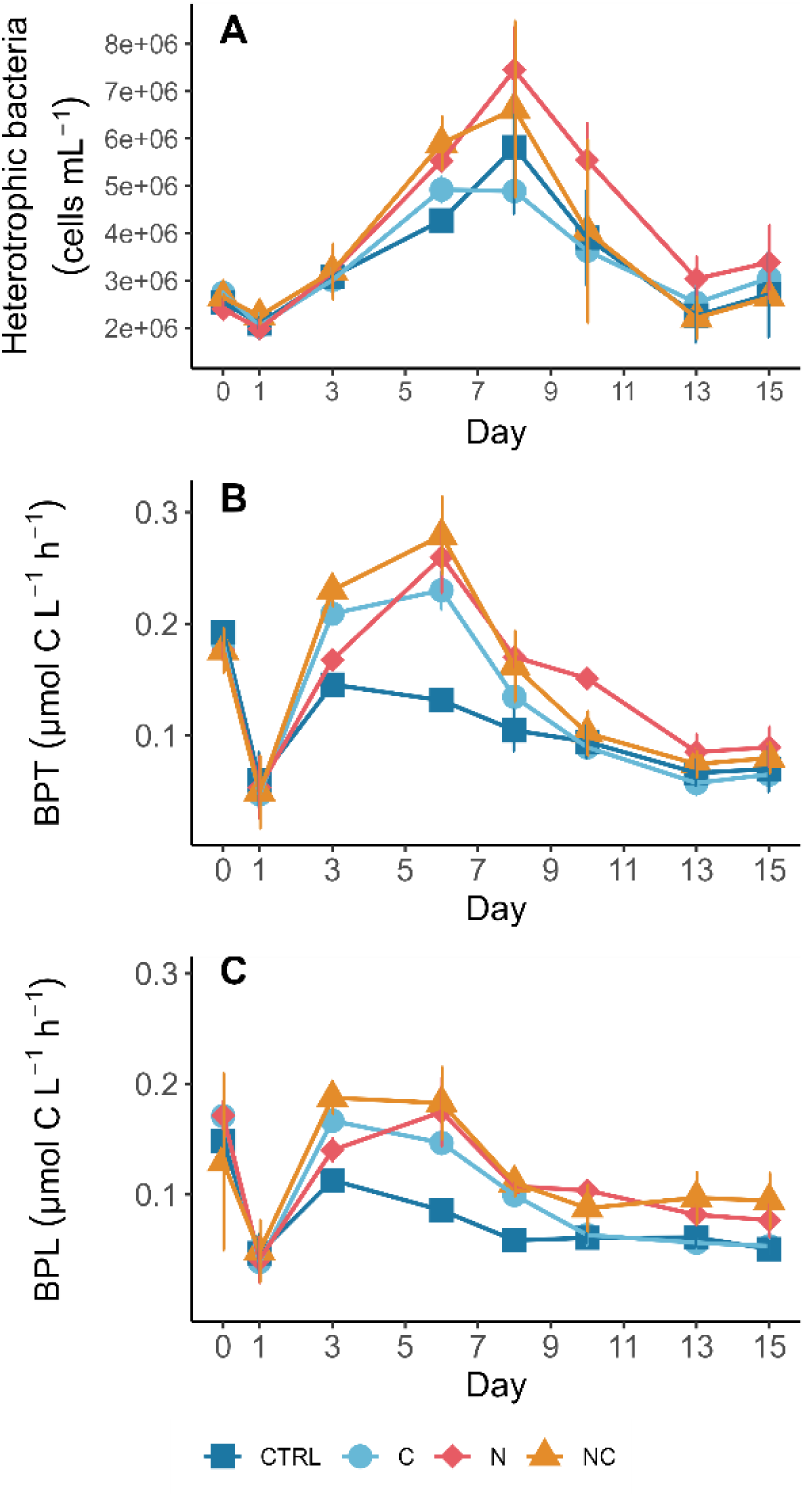
(A) Abundance of heterotrophic bacteria and bacterial production measured as (B) thymidine incorporation (BPT) and (C) leucine incorporation (BPL) over time. Error bars represent standard deviations of triplicate mesocosm bags.

### Development and drivers of extracellular enzyme activities

Similar to bacterial production (Fig. 3. B,C), the temporal development of bacterial abundance followed closely the activities of AGase, BGase, and LAPase in the fraction 0.2–3 µm with a lag of two days (Fig. 4). LAPase had the highest hydrolytic rates compared to other enzymes. During the first half of the experiment, activities of AGase, BGase, and LAPase were predominant in the bacterial (0.2–3 µm) fraction (Fig. 4) and the cell-specific AGase, BGase and LAPase showed two distinctive peaks during the experiment: the first one on days 3–5 and the second one after day 10 (Fig. 5). Cell-specific BPT correlated positively with cell-specific AGase, BGase, and LAPase activities (p = 0.0082, p = 0.001, and p = 0.0003, respectively, Table S1). These activities were also positively affected by temperature (Table S1). In addition, peptidolytic and glycolytic activities in fraction 0.2–3 µm showed negative responses to chlorophyll-*a* and DOC concentrations (Table S1). Conversely, there was no temporal congruence between APase activity in the 0.2–3 µm fraction and bacterial abundance and significant explanatory variables included DIP and POP concentrations with a positive effect (p < 0.0001, Table S1) and less significant negative effects of chlorophyll-*a* (p = 0.0023, Table S1) and temperature (p = 0.0437, Table S1) on APase activity.

**Fig. 4.**
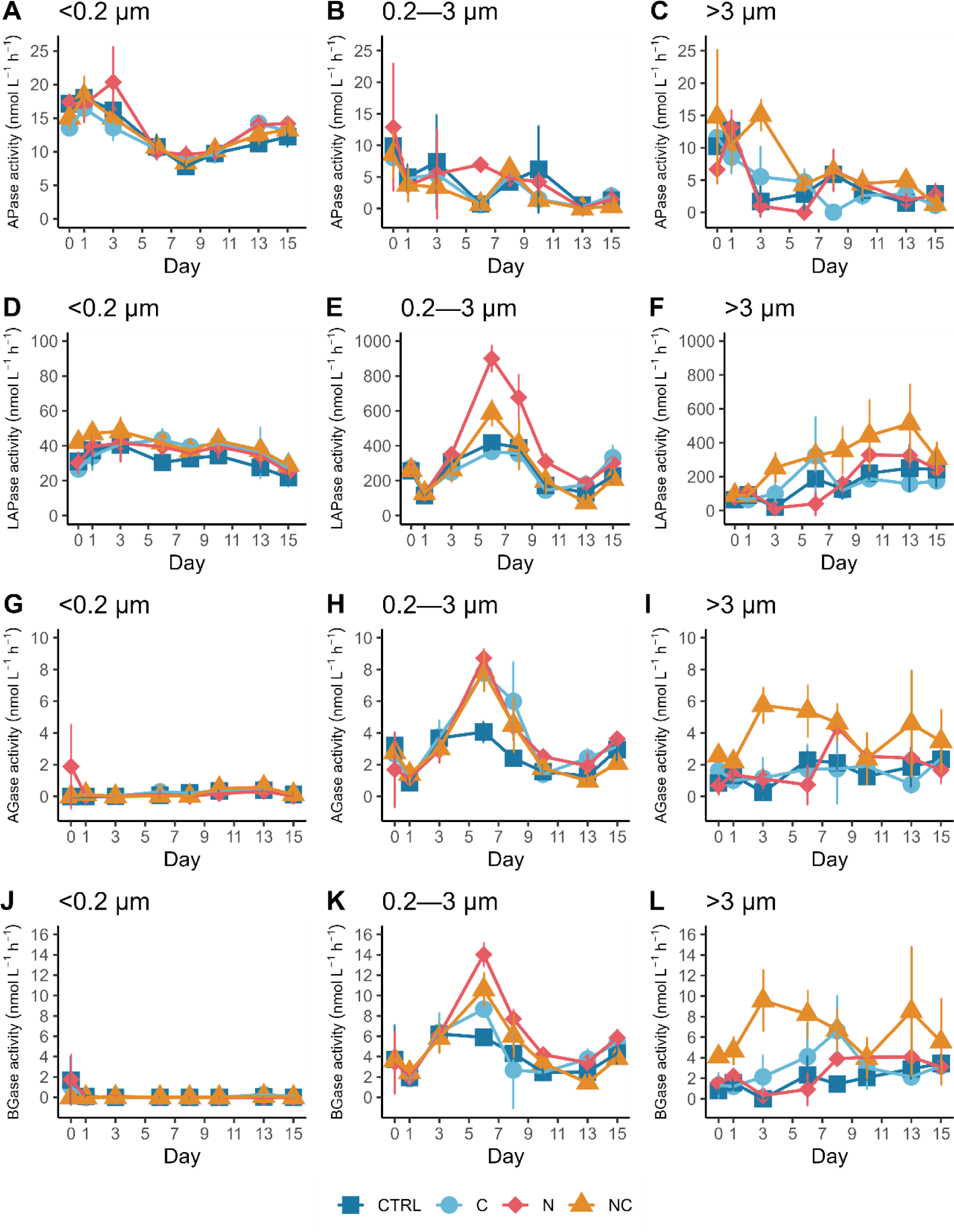
Potential hydrolytic activities (nmol L^-1^ h^-1^) of (A, B, C) alkaline phosphatase (APase), (D, E, F) leucine aminopeptidase (LAPase), (G, H, I) α-glucosidase (AGase), and (J, K, L) β-glucosidase (BGase) in three fractions (A, D, G, J) < 0.2 µm, (B, E, H, K) 0.2–3 µm, and (C, F, I, L) > 3 µm. Note that the scale for LAPase (D) is an order of magnitude lower than for the two other fractions (E and F). Error bars represent standard deviations of triplicate mesocosm bags. Non-visible error bars are smaller than the marker.

**Fig. 5.**
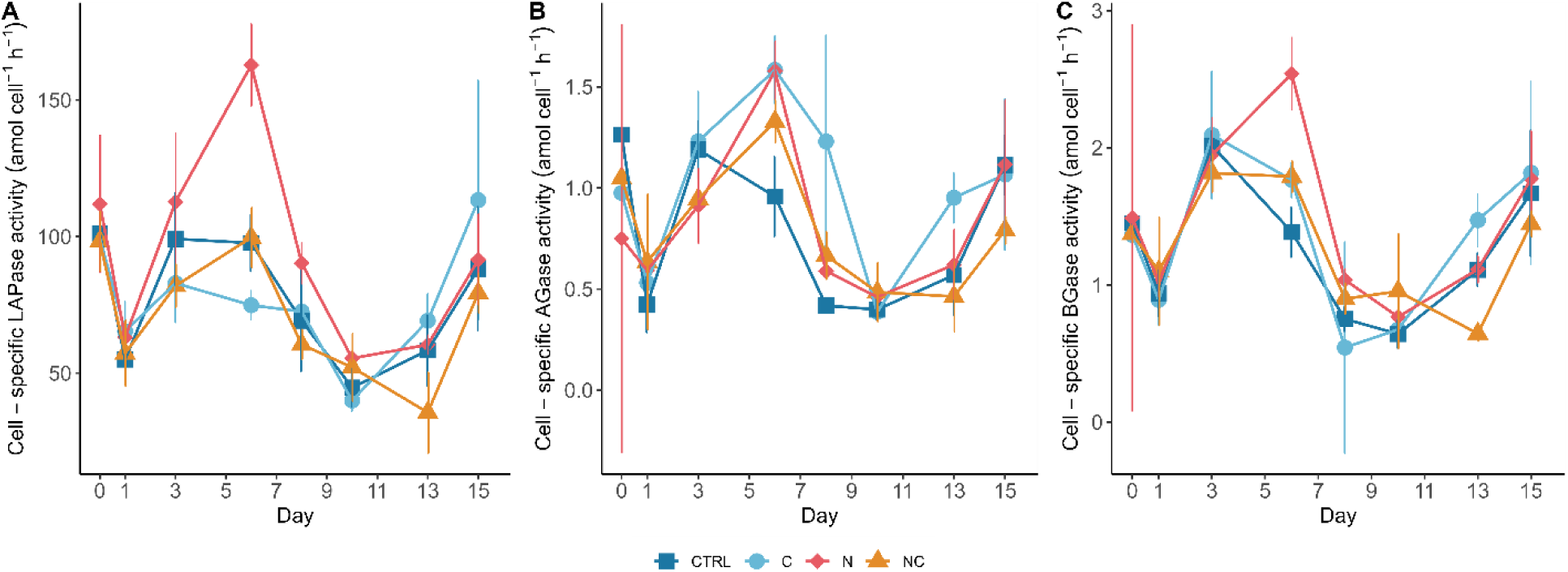
The development of cell-specific extracellular enzyme activities of (A) leucine-aminopeptidase (LAPase), (B) α-glucosidase (AGase), and (C) β-glucosidase (BGase) in the size fraction 0.2–3 µm. Error bars represent standard deviations of triplicate mesocosm bags.

The dissolved APase fraction accounted for the largest share of total APase activity throughout the experiment (on average 60.3 %) (Fig. S2). With a slight increase after the mid experiment, the dissolved fraction was the only fraction of APase activity that did not show a decreasing trend over time (Fig. 4). The dissolved APase activity was positively affected by temperature (p < 0.0001) and chlorophyll-*a* concentration (p < 0.0001). In contrast to high dissolved APase activities, activity of dissolved glucosidases was negligible (Fig. 4 and Fig. S2) and the dissolved LAPase activity was substantially lower compared to other fractions constituting 4–15 % of total LAPase activity. In addition, it showed a slight decreasing trend over time without any treatment effect (Fig. 4). Dissolved LAPase displayed a significant negative response to temperature (p < 0.0001, Table S1) and a positive relationship with NH_4_^+^ concentration (p < 0.0001, Table S1).

Peptidolytic rates in the >3 µm fraction increased over time with slightly higher LAPase rate in the NC-treatment (Fig. 4F), but the treatment effect was insignificant (p = 0.0922). LAPase activity in the >3 µm fraction showed a significant negative response to NH_4_^+^ concentration (p = 0.0133), temperature (p = 0.0335), and POC:PON ratio (p = 0.0472) (Table S1). Glycolytic activities in the >3 µm fraction had a clear peak in the NC-treatment on experiment day 3 and a smaller peak on day 13 (Fig. 4). For the AGase activities in fraction >3 µm, the NC-treatment displayed significantly higher rates compared to the control (p = 0.0259, Table S1), the N-treatment (p =0.0216, Table S2), and the C-treatment (p = 0.0067, Table S2), while BGase activity in the NC-treatment was significantly higher in comparison to the control (p = 0.0177, Table S2). None of the explanatory variables significantly affected BGase in the >3 µm fraction, but on AGase temperature had a slight negative effect (p = 0.0269) and DOC concentration a slight positive effect (p = 0.0316). The two most significant variables affecting APase activity in the >3 µm fraction were POP concentration and POC:POP ratio with a positive correlation (p < 0.0001, p = 0.0028, respectively, Table S1).

### Bacterial community structure and functional predictions

Based on 16S rRNA gene sequences, the most abundant classes over the experiment were Acidimicrobiia, Actinobacteria, Alphaproteobacteria, Bacteroidia, Bacilli, Cyanobacteria, Gammaproteobacteria, Planctomycetes, and Verrucomicrobiae (Fig. 6). During the first half of the experiment, Actinobacteria was the dominating bacterial class with an abundance ranging between 39 % on day 0 and 36 % on day 3 across all mesocosms followed by a decrease in relative abundance of Alphaproteobacteria from 21 % on day 3 to 17 % on day 6 and a slight increase in relative abundance of Bacteroidia from 17 % on day 3 to 21 % on day 6. In the second half of the experiment, an increase in relative abundance was observed for the classes Cyanobacteria ranging between 10 % on day 8 and 22 % on day 15, Gammaproteobacteria between 16 % on day 8, and 18 % on day 15, and Planctomycetes from 3 % on day 10 to 5 % on day 15. Simultaneously, a decrease in abundance of Actinobacteria from 31 % to 10 % between days 8 and 15 was observed. ASVs contributions to the predicted metabolic activities of LAPase, AGase, BGase and APase were investigated with the aim to describe the interplay between the most abundant bacterial taxa and the predicted metabolic processes within the community. Based on taxonomic affiliation, fifteen ASVs were predicted to encode LAPase, AGase, and BGase whereas only six ASVs were predicted to encode APase. Nearly the same ASVs contributed to the predicted LAPase, AGase, and BGase activities, except for ASV1 classified as *Cyanobium gracile* PCC 6307 and ASV3 and ASV6 classified as *Candidatus* Limnoluna that only contributed to the predicted LAPase and AGase activities. The ASVs contributing to the predicted activities changed consistently with the succession of the bacterial community with a clear transition of the functional composition on day 8 of the experiment (Table 1). In general, ASVs classified as *Sporichthyaceae* (Actinobacteria) and *Pseudorhodobacter* (Alphaproteobacteria) contributed to the predicted LAPase, AGase, and BGase activities during the first half of the experiment (days 1–6) when the measured activities of these enzymes increased and peaked (Fig. 4). In addition, *Candidatus* Limnoluna (ASV3 and ASV6, Actinobacteria) contributed most to the predicted AGase activity on days 1–6. Contribution of *C. gracile* PCC 6307 (Cyanobacteria) to the total predicted LAPase and BGase activities increased on days 13 and 15 (up to 16 % of the community-wide function abundance). The groups contributing to the community-wide predicted activities at the end of the experiment differed from the ones present at the beginning, although they also included representatives from classes Actinobacteria and Alphaproteobacteria (Table 1). Furthermore, their contribution to the community-wide predicted activity was generally lower than the ones predicted for other taxa at the start of the experiment. *Pseudohongiella* (Gammaproteobacteria) and *Isosphaeraceae* (Planctomycetes) contributed to the predicted activities only during the last sampling days.

**Fig. 6.**
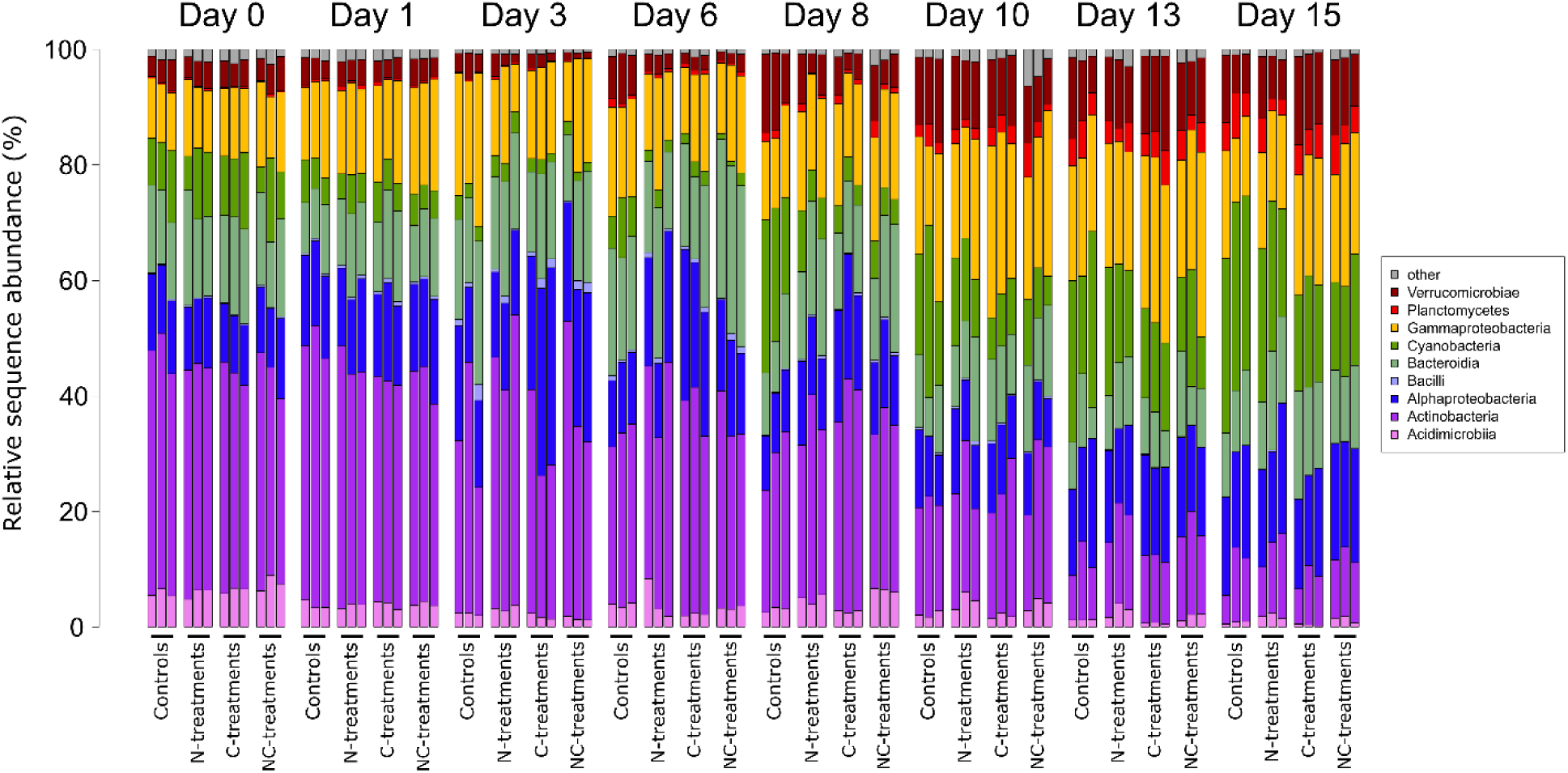
Temporal development of the 10 most abundant classes of the bacterial community.

**Table 1.**
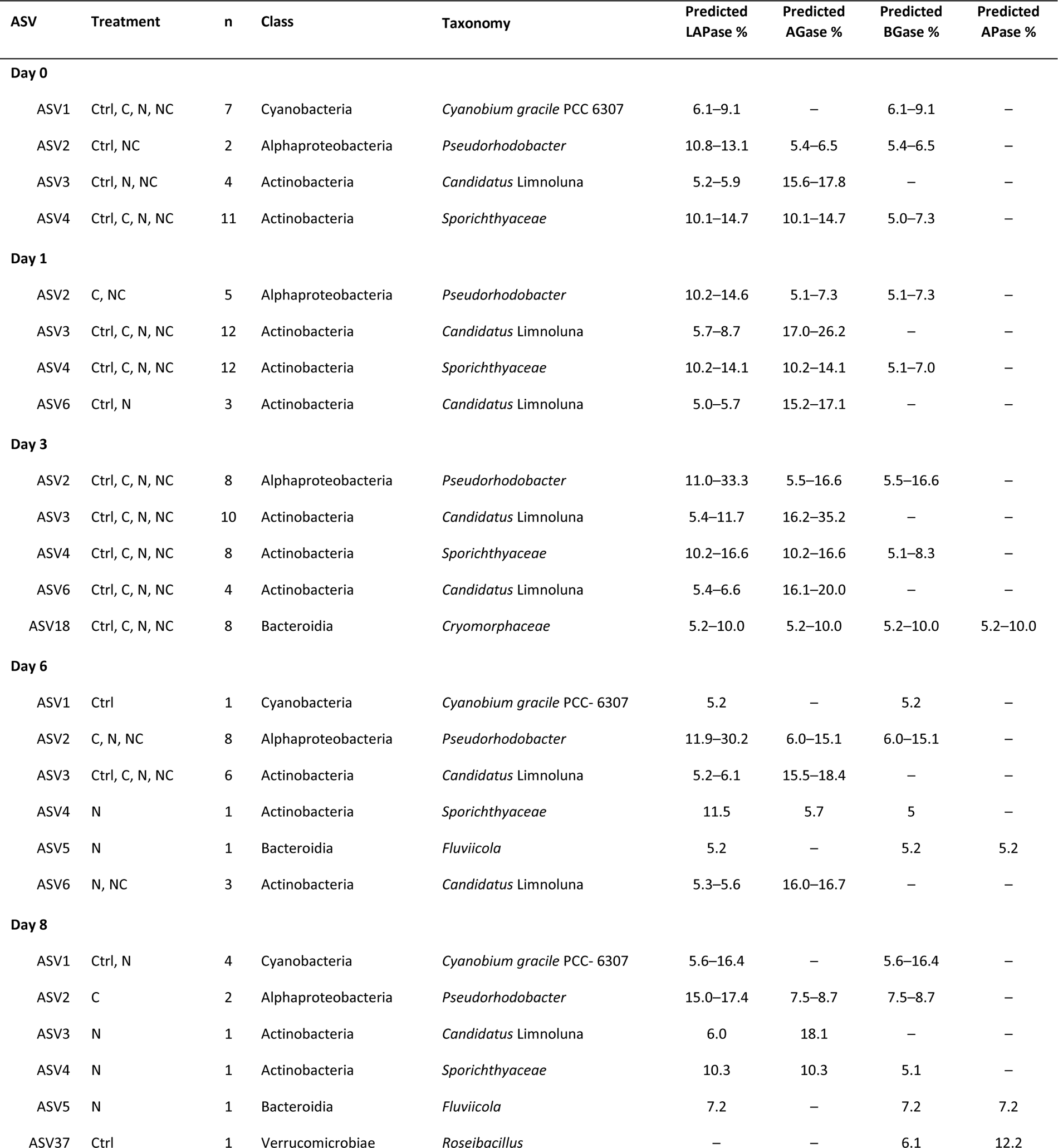
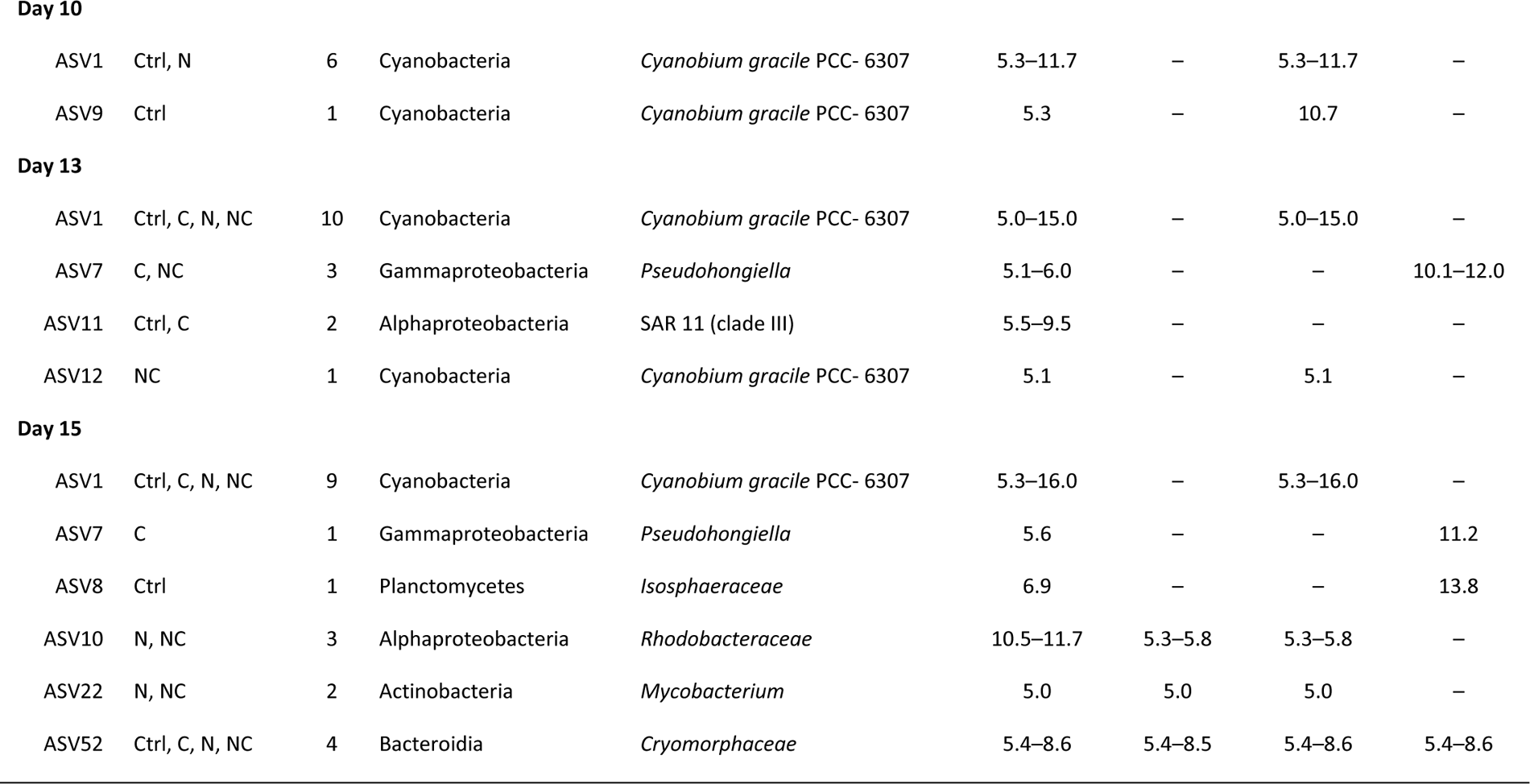
List of ASVs that contributed to the predicted enzymatic activities (expressed as relative abundance of metabolic functions) throughout the experiment with ranges between minimum and maximum values. Column n indicates the number of mesocosms where the respective taxon occurred. Column Taxonomy indicates the lowest classified taxonomic level for each ASV. The symbol – indicates a lower contribution than 5 % of an ASV to the predicted enzymatic activity.

| ASV | Treatment | n | Class | Taxonomy | Predicted<br>LAPase % | Predicted<br>AGase % | Predicted<br>BGase % | Predicted<br>APase % |
| --- | --- | --- | --- | --- | --- | --- | --- | --- |
| **Day 0** |  |  |  |  |  |  |  |  |
| ASV1 | Ctrl, C, N, NC | 7 | Cyanobacteria | *Cyanobium gracile* PCC 6307 | 6.1–9.1 | – | 6.1–9.1 | – |
| ASV2 | Ctrl, NC | 2 | Alphaproteobacteria | *Pseudorhodobacter* | 10.8–13.1 | 5.4–6.5 | 5.4–6.5 | – |
| ASV3 | Ctrl, N, NC | 4 | Actinobacteria | *Candidatus* Limnoluna | 5.2–5.9 | 15.6–17.8 | – | – |
| ASV4 | Ctrl, C, N, NC | 11 | Actinobacteria | *Sporichthyaceae* | 10.1–14.7 | 10.1–14.7 | 5.0–7.3 | – |
| **Day 1** |  |  |  |  |  |  |  |  |
| ASV2 | C, NC | 5 | Alphaproteobacteria | *Pseudorhodobacter* | 10.2–14.6 | 5.1–7.3 | 5.1–7.3 | – |
| ASV3 | Ctrl, C, N, NC | 12 | Actinobacteria | *Candidatus* Limnoluna | 5.7–8.7 | 17.0–26.2 | – | – |
| ASV4 | Ctrl, C, N, NC | 12 | Actinobacteria | *Sporichthyaceae* | 10.2–14.1 | 10.2–14.1 | 5.1–7.0 | – |
| ASV6 | Ctrl, N | 3 | Actinobacteria | *Candidatus* Limnoluna | 5.0–5.7 | 15.2–17.1 | – | – |
| **Day 3** |  |  |  |  |  |  |  |  |
| ASV2 | Ctrl, C, N, NC | 8 | Alphaproteobacteria | *Pseudorhodobacter* | 11.0–33.3 | 5.5–16.6 | 5.5–16.6 | – |
| ASV3 | Ctrl, C, N, NC | 10 | Actinobacteria | *Candidatus* Limnoluna | 5.4–11.7 | 16.2–35.2 | – | – |
| ASV4 | Ctrl, C, N, NC | 8 | Actinobacteria | *Sporichthyaceae* | 10.2–16.6 | 10.2–16.6 | 5.1–8.3 | – |
| ASV6 | Ctrl, C, N, NC | 4 | Actinobacteria | *Candidatus* Limnoluna | 5.4–6.6 | 16.1–20.0 | – | – |
| ASV18 | Ctrl, C, N, NC | 8 | Bacteroidia | *Cryomorphaceae* | 5.2–10.0 | 5.2–10.0 | 5.2–10.0 | 5.2–10.0 |
| **Day 6** |  |  |  |  |  |  |  |  |
| ASV1 | Ctrl | 1 | Cyanobacteria | *Cyanobium gracile* PCC- 6307 | 5.2 | – | 5.2 | – |
| ASV2 | C, N, NC | 8 | Alphaproteobacteria | *Pseudorhodobacter* | 11.9–30.2 | 6.0–15.1 | 6.0–15.1 | – |
| ASV3 | Ctrl, C, N, NC | 6 | Actinobacteria | *Candidatus* Limnoluna | 5.2–6.1 | 15.5–18.4 | – | – |
| ASV4 | N | 1 | Actinobacteria | *Sporichthyaceae* | 11.5 | 5.7 | 5 | – |
| ASV5 | N | 1 | Bacteroidia | *Fluviicola* | 5.2 | – | 5.2 | 5.2 |
| ASV6 | N, NC | 3 | Actinobacteria | *Candidatus* Limnoluna | 5.3–5.6 | 16.0–16.7 | – | – |
| **Day 8** |  |  |  |  |  |  |  |  |
| ASV1 | Ctrl, N | 4 | Cyanobacteria | *Cyanobium gracile* PCC- 6307 | 5.6–16.4 | – | 5.6–16.4 | – |
| ASV2 | C | 2 | Alphaproteobacteria | *Pseudorhodobacter* | 15.0–17.4 | 7.5–8.7 | 7.5–8.7 | – |
| ASV3 | N | 1 | Actinobacteria | *Candidatus* Limnoluna | 6.0 | 18.1 | – | – |
| ASV4 | N | 1 | Actinobacteria | *Sporichthyaceae* | 10.3 | 10.3 | 5.1 | – |
| ASV5 | N | 1 | Bacteroidia | *Fluviicola* | 7.2 | – | 7.2 | 7.2 |
| ASV37 | Ctrl | 1 | Verrucomicrobiae | *Roseibacillus* | – | – | 6.1 | 12.2 |
| **Day 10** |  |  |  |  |  |  |  |  |
| ASV1 | Ctrl, N | 6 | Cyanobacteria | *Cyanobium gracile* PCC- 6307 | 5.3–11.7 | – | 5.3–11.7 | – |
| ASV9 | Ctrl | 1 | Cyanobacteria | *Cyanobium gracile* PCC- 6307 | 5.3 | – | 10.7 | – |
| **Day 13** |  |  |  |  |  |  |  |  |
| ASV1 | Ctrl, C, N, NC | 10 | Cyanobacteria | *Cyanobium gracile* PCC- 6307 | 5.0–15.0 | – | 5.0–15.0 | – |
| ASV7 | C, NC | 3 | Gammaproteobacteria | *Pseudohongiella* | 5.1–6.0 | – | – | 10.1–12.0 |
| ASV11 | Ctrl, C | 2 | Alphaproteobacteria | SAR 11 (clade III) | 5.5–9.5 | – | – | – |
| ASV12 | NC | 1 | Cyanobacteria | *Cyanobium gracile* PCC- 6307 | 5.1 | – | 5.1 | – |
| **Day 15** |  |  |  |  |  |  |  |  |
| ASV1 | Ctrl, C, N, NC | 9 | Cyanobacteria | *Cyanobium gracile* PCC- 6307 | 5.3–16.0 | – | 5.3–16.0 | – |
| ASV7 | C | 1 | Gammaproteobacteria | *Pseudohongiella* | 5.6 | – | – | 11.2 |
| ASV8 | Ctrl | 1 | Planctomycetes | *Isosphaeraceae* | 6.9 | – | – | 13.8 |
| ASV10 | N, NC | 3 | Alphaproteobacteria | *Rhodobacteraceae* | 10.5–11.7 | 5.3–5.8 | 5.3–5.8 | – |
| ASV22 | N, NC | 2 | Actinobacteria | *Mycobacterium* | 5.0 | 5.0 | 5.0 | – |
| ASV52 | Ctrl, C, N, NC | 4 | Bacteroidia | *Cryomorphaceae* | 5.4–8.6 | 5.4–8.5 | 5.4–8.6 | 5.4–8.6 |

## Discussion

The aim of this study was to investigate the possible drivers of EEAs and the degradation potential of OM which is crucial in determining the utilization of the excess phosphate pool and export of OM to the sea floor following the northern Baltic spring bloom. In addition, measured EEAs were compared with predicted functions identified with their corresponding enzymatic activities based on 16S rRNA gene sequences of the bacterial community.

### Nitrogen and carbon deficient heterotrophic bacteria

The role of heterotrophic bacteria in excess phosphate utilization was of specific interest due to their generally high cellular phosphorus content and low C:P ratio (Chrzanowski and Kyle 1996, Heldal et al. 1996). Their abundance peak around day 8 occurred after the peaks of bacteria-specific glycolytic and peptidolytic activities and bacterial production, which reached relatively high rates compared to recent studies in the Baltic Sea (Bunse et al. 2019, Camarena-Gómez et al. 2020, Zhao et al. 2023). AGase, BGase, and LAPase activities in the 0.2–3 µm fraction showed a similar temporal pattern and were clearly related to the bacterioplankton development, whereas APase was not. In line with this finding, temperature positively influenced cell-specific AGase, BGase, and LAPase activities while displaying a negative effect on APase activity in the 0.2–3 µm fraction (Table S1). A correlation between BGase and LAPase has been observed before (Grossart et al. 2006, Baltar et al. 2016, Cai et al. 2022), suggesting that such a relationship is common in coastal areas. The most abundant bacterial taxa were responsible for the contribution of both peptidolytic and glycolytic predicted activities (Table 1), which further demonstrates that degradation of N and C were tightly coupled also at the organismic level.

Cell-specific BPT showed a significant response with cell-specific LAPase, BGase and AGase activities demonstrating a tight coupling between substrate degradation and uptake. These findings suggest that growth of heterotrophic bacteria, especially of taxa belonging to the class Actinobacteria, was driven by hydrolysis of N- and C-rich OM indicating inorganic nitrogen and labile carbon deficiency, that was dealt with by utilizing available high molecular weight substrates that are typically associated with senescent phytoplankton and termination of phytoplankton blooms (Chróst 1991, Middelboe et al. 1995). Members of Actinobacteria are known to contribute to N and C cycling by degrading chitin that can derive from phyto- and zooplankton (Tada and Grossart 2014). Negative response of cell-specific glycolytic and peptidolytic activities to either or both chlorophyll-*a* and DOC concentrations further support this interpretation (Table S1). Consistently with our results, Actinobacteria are known to occur in post-bloom conditions especially in the less saline northern Baltic Sea (Hugerth et al. 2015, Rieck et al. 2015, Bunse et al. 2016) and to incorporate thymidine (Piwosz et al. 2013).

Bacterial LAPase rates in our study were several times higher compared to those measured in most previous studies, but in line with bulk LAPase activities in the Peruvian coastal upwelling system (200–800 nmol L^-1^ h^-1^; Spilling et al. 2023), which is an oxygen minimum zone with a low DIN:DIP ratio. In the northern Baltic, post-spring-bloom low DIN:DIP conditions, high N_2_-fixation and degradation of N-rich organic compounds can potentially support the drawdown of excess phosphate. During our experiment, regenerated production likely prevailed due to the low abundance of large N_2_-fixing filamentous cyanobacteria (Santoro et al., in prep.). Therefore, we assume that excess phosphate might enhance the hydrolytic rates of bacterial peptidolytic and glycolytic enzymes, but probably only in combination with organic nitrogen substrates (Nausch 2000). The added DIN seemed to be transformed quickly into DON, which was then utilized presumably by the free-living bacteria showing increasing LAPase rates during the first half of the experiment (Fig. 4E). Increasing DON concentration during the second half indicates accumulation of hydrolysis resistant DON of bacterial origin (Ogawa et al. 2001).

Although glucose addition enhanced bacterial production rates, C- and NC-treatments did not seem to give any advantage for heterotrophic bacteria as the effect was similar to that in the control (Fig. 3). This notion, together with relatively high glycolytic activities in the C-amendment, indicate that glucose addition was insufficient for heterotrophic bacteria and the added glucose might have been used to primarily enhance respiration (Guillemette et al. 2016), which is also supported by the lack of an obvious C-treatment effect on bacterial abundance (Fig. 3A). High LAPase activity might have alleviated the carbon demand of heterotrophic bacteria as the degradation of peptides by LAPase is not only utilized for acquiring N but also for providing C (Ylla et al. 2012). During post-spring-bloom conditions, the bacterial community could be well adapted to utilizing algal-derived polymeric substrates (Eigemann et al. 2022, Villena-Alemany et al. 2023), which could also reduce the effect of glucose addition on bacterial abundance. ASVs belonging to the class Bacteroidia, typically linked to the degradation of high molecular weight OM in the ocean (Kirchman 2002), contributed to the predicted glycolytic activities in our experiment. However, their contribution to the total predicted BGase and AGase activities ranged only between 5.2 and 10.0 %, although their abundance was relatively high (∼21 %) on day 6. *Rhodobacteraceae*, a group in the order Rhodobacterales known to utilize recalcitrant OM (Allers et al. 2007), appeared at the end of the experiment. This might indicate reduced availability of labile energy sources presumably limiting both abundance and activity of heterotrophic bacteria, reducing their ability to compete for inorganic nutrients and providing advantage for picocyanobacteria at the end of the experiment. In support of this, the decline of BPT and BPT:BPL ratio to the initial level at the end of the experiment indicated suboptimal conditions for bacterial growth (Fig. 3C).

### Fate of excess phosphate

Although the excess phosphate pool decreased in all treatments, it was not depleted during our experiment contradicting findings of a previous longer study in the Tvärminne archipelago (Vanharanta and Spilling 2023). APase activity has typically been used as an indicator of physiological P limitation of microbes (Nausch and Nausch 2004, Mahaffey et al. 2014, Baltar et al. 2016), but it can also be used for carbon acquisition in phosphate replete systems (Hoppe 2003). The hydrolytic rates of bulk APase measured in our study under phosphate replete conditions (12.3–44.4 nmol L^-1^ h^-1^) were comparable to the activities measured in May–June in the western Baltic Proper (0–31.9 nmol L^-1^ h^-1^) that were associated with P limitation (Baltar et al. 2016). It is plausible that the hydrolytic rates measured in our study had been relicts from P-limiting conditions before the start of the experiment as indicated by DIP concentration below the threshold of 0.03 µmol L^-1^ known to induce APase production (Mahaffey et al. 2014) and the decreasing trend of APase in the two larger fractions over time. This finding demonstrates the long extracellular lifetime of these enzymes (Thomson et al. 2019). In other P-limited aquatic environments, however, many-fold higher APase activities have been reported, e.g. Lake Constance (115 nmol L^-1^ h^-1^; Grossart and Simon 1998), the southern Baltic Sea (543 nmol L^-1^ h^-1^; Nausch et al. 1998), Florida Bay (1220 nmol L^-1^ h^-1^; Koch et al. 2009) and the Adriatic Sea (2916 nmol L^-1^ h^-1^; Ivančić et al. 2016) demonstrating the difficulty in relating EEAs to nutrient deficiency of bacterial communities and to compare enzymatic hydrolytic rates across different aquatic ecosystems.

We detected an increase of ∼0.2 µmol of DOP L^-1^ during the first half of the experiment, which was mineralized between days 6 and 15 although phosphate was still available. It is probable that dissolved APase, constituting the majority of the total APase activity (44 %–90 %, Fig. S2), contributed to the observed decrease in DOP concentration after mid experiment and thereby to excess phosphate concentrations. The possible source for dissolved APase during the second half of the experiment are carbon limited heterotrophic bacteria known to secrete APase into the solution (Luo et al. 2009). This is corroborated by the metabolic prediction results that showed an increasing contribution to the predicted APase activity of bacterial ASVs after day 8 (Table 1). In particular, ASVs in classes Verrucomicrobiae and Planctomycetes seemed to require more phosphorus compared to other ASVs (Table 1). The highly significant correlation between chlorophyll-a and dissolved APase does probably not imply causation as phytoplankton are not expected to produce APase under phosphate replete conditions. Given the absence of correlation between APase activity in the size range 0.2–3 µm and bacterial abundance and the negative correlation with chlorophyll-*a* and temperature (Table S1), it seems reasonable that most APase activity in this size fraction was not related to free-living bacteria or small phytoplankton, but probably attributed to dissolved enzymes absorbed to small abiotic surfaces, although the true origin of APase cannot be confirmed. Furthermore, the positive correlation between APase activity in the fraction >3 µm and POP concentration and the POC:POP ratio (Table S1) could indicate enzymatic APase activity from particle attached bacteria.

### Particle-associated and dissolved enzymatic activities and their implication for downward fluxes

In the NC-treatment, significantly higher rates of AGase, and BGase activities in the >3 µm fraction likely resulted from a higher biovolume (Fig. S1A) serving as surfaces for microbial colonization and enhanced EEAs (Simon et al. 2002). The ability of particle-attached bacteria to exploit a wider variety of polymeric C-substrates compared to their free-living counterparts (Lyons and Dobbs 2012) has been shown to result in higher enzymatic activities on aggregates than in the surrounding water (e.g. Grossart et al. 2003). However, our results contradict with several studies that showed higher EEAs of particle-attached bacteria compared to free-living ones (Karner and Herndl 1992, Smith et al 1992, Grossart et al. 2007), as activities of LAPase, AGase and BGase in the size fraction > 3 µm were comparable or generally lower than those measured in the 0.2–3 µm fraction (Fig. 4).

We did not count the abundance of particle-attached bacteria. However, it is possible that, per sample volume, numbers of free-living bacteria exceeded those attached to aggregates due to high sedimentation losses observed as decreasing POM concentrations during the first days of the experiment, which was likely the consequence of nano- and microphytoplankton sinking out from the surface and being replaced by picophytoplankton after nitrate depletion as observed before in Hornick et al. (2017) (Spilling et al. REF). Bacterial growth on aggregates can also be balanced by intensive grazing of protozoans (Ploug and Grossart 2000) or even mesozooplankton (e.g. Steinberg et al. 2008). The detected increase in the abundance of copepods towards the end of the experiment (Spilling et al. REF) might have contributed to decreasing POM, which is further supported by our observations of suspected large, fragmented copepod fecal pellets (Fig. S3).

It is also likely that the two bacterial fractions differ phylogenetically and functionally from each other (Lyons and Dobbs 2012, Mohit et al. 2014, Rieck et al. 2015), which could lead to distinct enzymatic activities between both fractions (Azúa et al. 2003). Temporal succession of ASVs contributing to the predicted activities seemed to reflect a change from bacteria in the free-living to the particle-attached fraction. For instance, several ASVs classified as *Pseudorhodobacter*, *Candidatus* Limnoluna, and *Sporichthyaceae*, were predicted to contribute to total enzymatic activities only during the first half of the experiment. Of these, *Sporichthyaceae* (Actinobacteria) has been associated with a free-living lifestyle (Bashenkhaeva et al. 2020). By contrast, *Cryomorphaceae* (Bacteroidia) and *Mycobacterium* (Actinobacteria) have been associated with a particle-attached lifestyle (Delmont et al. 2014, Reintjes et al. 2023, Allgaier et al. 2007), and from these, contribution of *Mycobacterium* to the community-wide predicted LAPase activity was observed only at the end of the experiment.

For surface-attached bacteria, it might be profitable to release hydrolytic enzymes into solution due to high substrate concentrations on particles (Vetter et al. 1998), in which case we would measure the part of enzymatic activity that escapes aggregates in the dissolved fraction. Regarding export fluxes, OM quality is considered an important characteristic for microbial uptake and metabolism (Boyd and Trull 2007). The observed POC:PON ratio was above the Redfield ratio throughout the entire experiment (Fig. 2J), consequently the contribution of detritus to total biovolume was relatively high (Fig. S1B). Regardless, peptidolytic and glycolytic EEAs shifted from being predominantly associated with free-living bacteria towards the >3 µm fraction at the end of the experiment. This indicates intense bacterial particle colonization and simultaneous succession of the bacterial community as a consequence of temporal changes in available substrates. This would suggest attenuation of carbon flux to the seafloor. Higher N degradation relative to C also in the >3 µm size fraction implies that POM lost N faster compared to C thereby increasing the C:N ratio of potentially sinking material (Smith et al. 1992, Grossart and Ploug 2001).

## Conclusions

Excess phosphate was not depleted during the experiment, and we found a relatively high potential of dissolved APase activity to play a substantial role in sustaining the surplus phosphate pool. These activities probably derived from microbial activity during phosphate limited conditions prior to the experiment and from carbon limited heterotrophic bacteria at the end of the experiment. The computational approach predicting functional activities via 16S rRNA gene sequences suggested that taxa belonging to Actinobacteria and Alphaproteobacteria were potentially important producers of peptidolytic and glycolytic enzymes during the bacterioplankton bloom development when accessible substrates were still available at the beginning of the mesocosm experiment. Our findings indicate N- and C-limitation of heterotrophic bacteria with a faster degradation rate of peptides compared to polysaccharides, which implies a higher dissolution rate of N-compared to C-rich OM (Grossart et al. 2000). Low organic energy supply for heterotrophic bacteria seemed to make picocyanobacteria superior competitors for inorganic nutrients towards the end of the experiment, and probably induced succession of heterotrophs towards particle colonizing taxa with a probable attenuation of carbon flux to the seafloor. Measuring vertical patterns of EEA in the northern Baltic Sea could elucidate the potential of OM degradation in different layers of the water column, a necessary approach towards a better understanding of export fluxes and nutrient cycles.

## Acknowledgements

We thank Joanna Norkko and Laura Kauppi for technical support, Jostein Solbakken and Göran Lundberg for assistance in setting up the mesocosms and Mervi Sjöblom, Jaana Koistinen and Kia Rautava for inorganic and organic nutrient analyses at the Tvärminne Zoological Station. We thank Onni Talas foundation interns; Emma Forss, Neea Hanström, Anni Leinonen and Marlena Grönqvist, for their help in the laboratory and field. We extend our thanks to Christiane Hassenrück for developing the bioinformatics tailored workflow partially used in this manuscript, her helpful suggestions, and valuable discussions regarding the metabolic prediction investigation. The study utilized the Finnish Environment Institute marine research infrastructure as a part of the national Finnish Marine Research Infrastructure (FINMARI) consortium.

## Data availability

Data from all measured parameters are available on PANGAEA (Vanharanta et al., DOI pending). The sequence data for this study have been deposited in the European Nucleotide Archive (ENA) at EMBL-EBI under accession number PRJEB72147 (https://www.ebi.ac.uk/ena/browser/view/PRJEB72147).

