## Supplemental material for "Microbial remineralization processes during post-spring-bloom excess phosphate in the northern Baltic Sea"

**Supplemental Text**

**Supplemental Methods S1: Supplementary description of methods - DNA extraction, bioinformatics processing of sequence reads for bacterial community composition and prediction of metabolic functions**

Brief description of slight modifications of DNA extraction protocol of Nercessian et al. (2005). Heat-sterilized zirconium beads of 3 sizes (0.1 mm: 0.15 g; 0.7 mm: 0.15 g, 1mm: 0.2 g), 750 µl of CTAB-buffer (final conc. 10 % w/v in 1.6 M sodium chloride solution and 240 mM potassium-phosphate-buffer pH 8.0), 75 µl SDS (final conc. 1% w/v),) 75 µl N-Lauroylsarcosin (final conc. 1 % w/v) and 10 µl of Proteinase K (1 mg ml^-1^) were added and incubated at 55°C for 30 min. Seven-hundred-fifty µl of phenol-chloroform-isoamylalcohol (25:24:1) were added and the samples were centrifuged at 16 000 g at 4°C for 10 min. Supernatant was transferred and 1 volume of chloroform was added, and centrifugation was repeated. 2 volumes of a polyethylene glycol (PEG) - sodium chloride (NaCl) solution (1.6 M NaCl - final concentration 9.4% w/v, PEG 6000 - final concentration 30% w/v) and 1 µl of linear polyacrylamide (GenElute LPA, Sigma) were added and incubated in followed by a dark for 2 hours at 4°C. Centrifugation at 17 000 g at 4°C for 90 min. Precipitated nucleic acids were washed in 1 ml ice-cold ethanol (conc. 70%), air dried and dissolved in 35 µl of dietdiethylpyrocarbonate (DEPC) water and stored at -20°C until sequencing. Taxonomic composition and phylogenetic diversity of the bacterial community was assessed after collecting samples for 16S marker gene analysis. Subsequently, community-wide metabolic functions were predicted. Around 100 000 raw sequenced read pairs per sample were generated yielding ~20 million for all the samples. The number of reads obtained from the sequenced samples over the experiment ranged from 14 241 to 130 485. The number of retained ASVs (amplicon sequence variants) processed in the downward analysis was 3587. Sequence data processing included the following steps: - general and primer clipping (the allowed mismatch to the primer sequences set with an error of 0.16); - screening with threshold ranges for filtering and trimming (truncation length of read 1 set at a minimum of 250 base pairs and a maximum of 280 base pairs; fragment maximum length set at 430 base pairs; truncation length of read 2 was calculated based on fragment maximum length subtracting the truncation length of read 1 plus 30; maximum expected error set at a minimum of 2 and at a maximum of 3); - ultimate filtering after a data evaluation step (truncation length of read 1 set at 260 base pairs with a maximum expected error of 3, truncation length of read 2 set at 200 base pairs with a maximum expected error of 3) and the maximum expected fragment length between 378 and 433 base pairs; - denoising and merging (loess error function set as default for learning the error rates; no fragments exceeding the insert size for merging were rescued by concatenation); - taxonomic classification (bootstrap cut-off set at a minimum of 70 and megablast with NCBI NT reference database and blasting with SILVA v138.1 reference database and PR2. v4.14.0 reference database for chloroplasts and mitochondria classification, subsequently removed from the downstream analysis); - taxonomic assignment and rank mapping via blasting with SILVA v138.1 reference database.

**Supplemental figures**

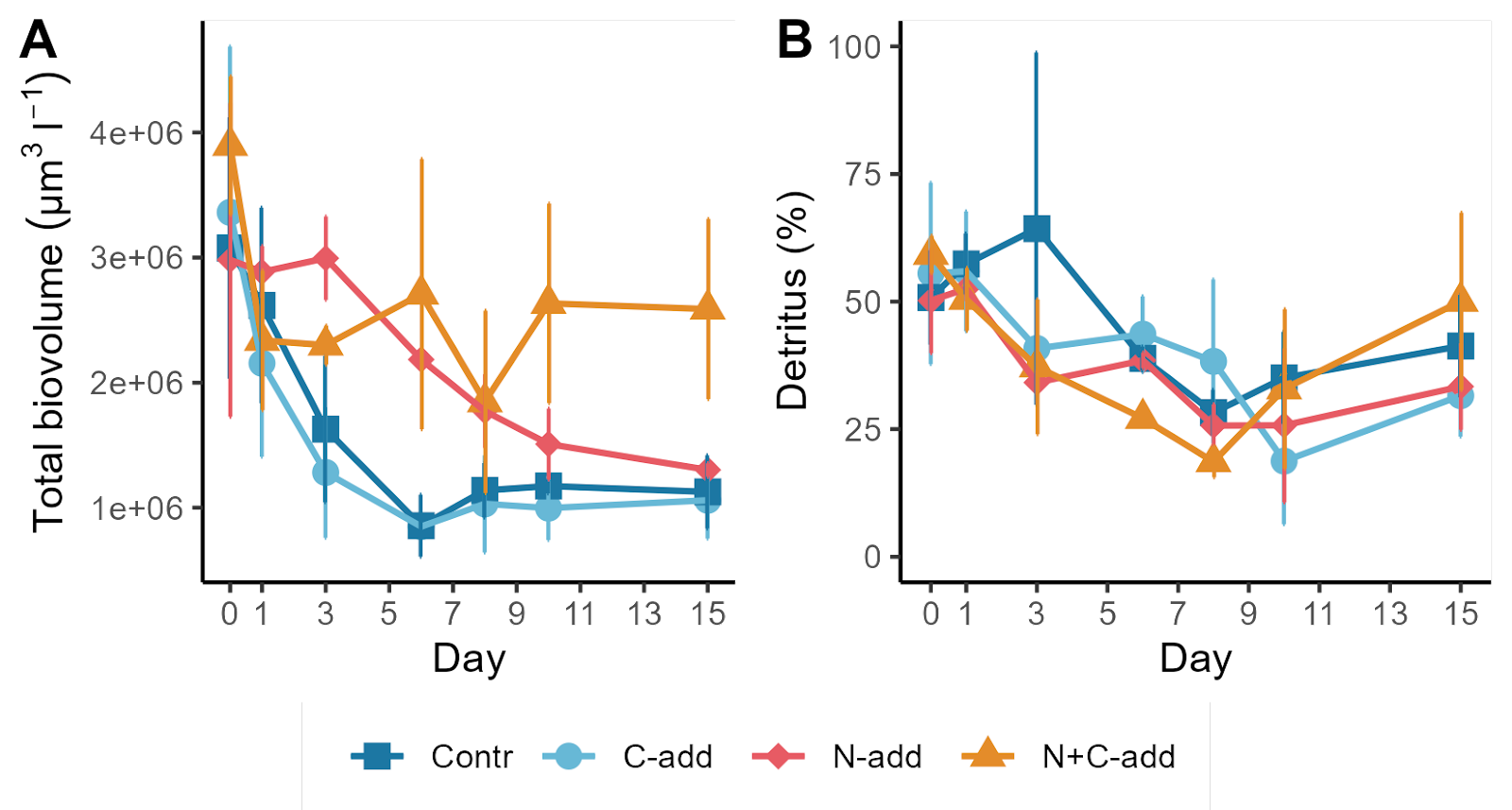
**Figure S1**. (A) Total biovolume in the mesocosms and (B) the contribution of detritus to the total organic matter biovolume over time. Error bars represent standard deviations of triplicate mesocosm bags.

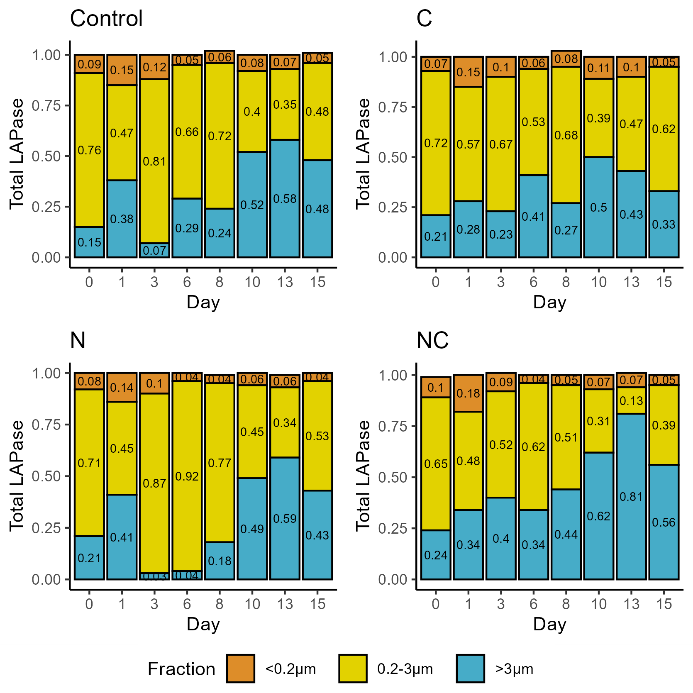

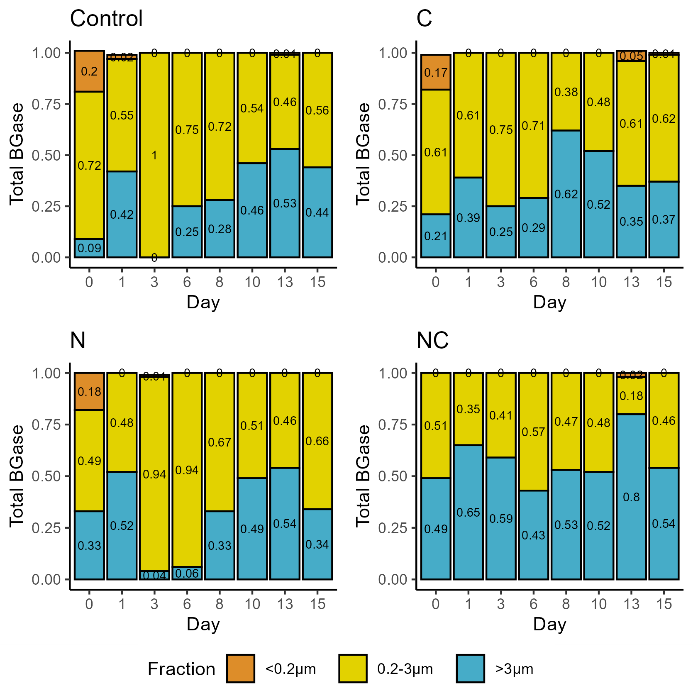

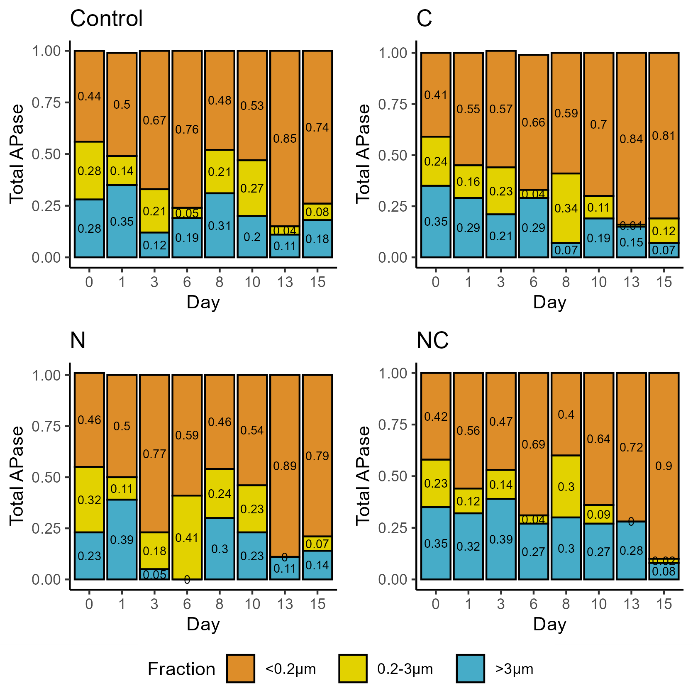

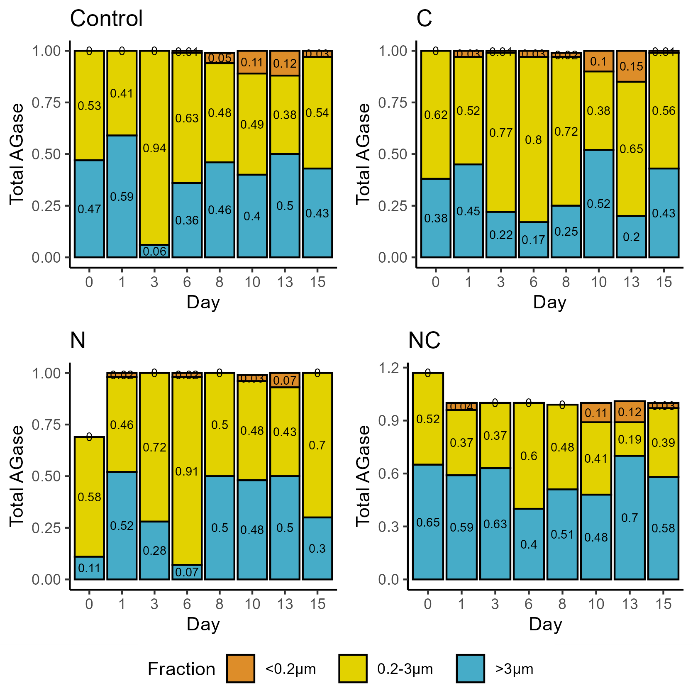

**Figure S2.** The development of relative contributions of different fractions in the four different treatments of the total extracellular enzymatic activities of LAPase, BGase, APase, and AGase.

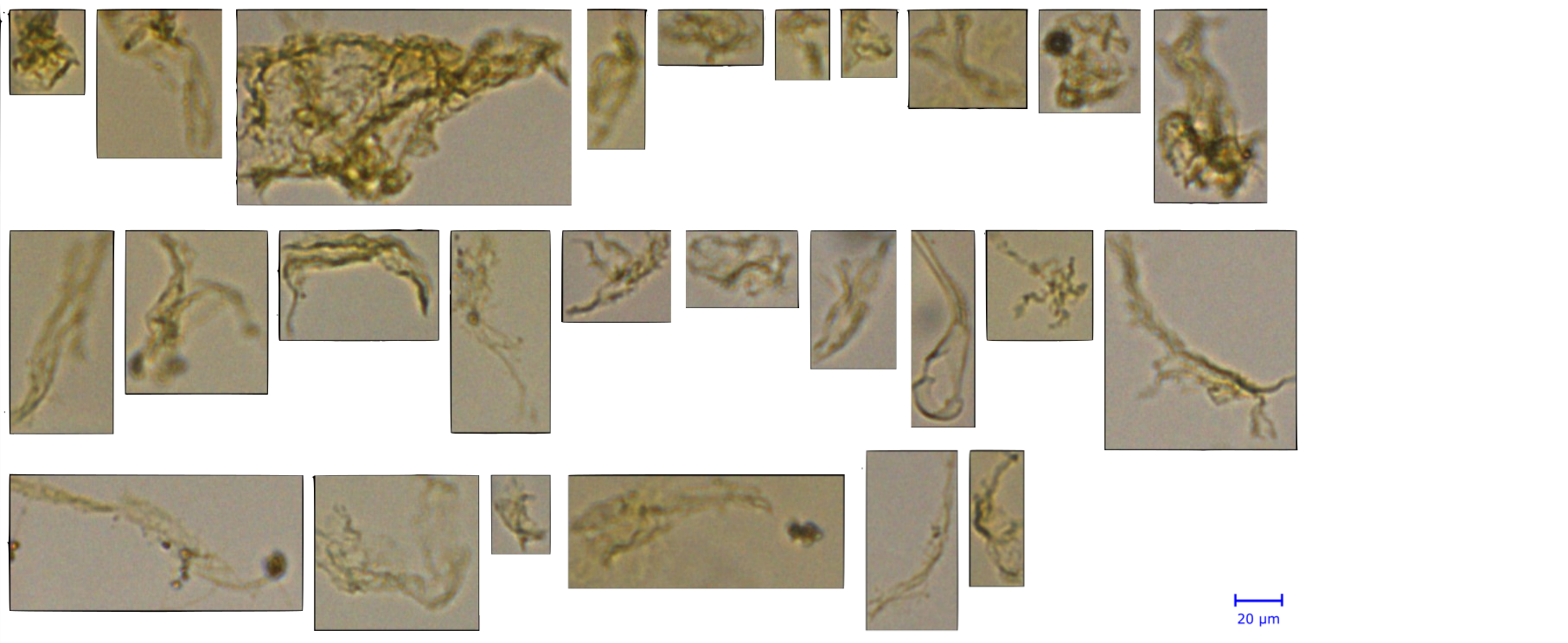

**Figure S3.** Fragmented copepod faecal pellets.

**Supplemental Tables**

**Table S1.** Summary outputs of the optimal mixed effects models showing the significant fixed effects for enzymatic activities in different size fractions. For AGase, BGase and LAPase in size fraction 0.2–3 µm, cell-specific enzymatic activity was used as response variable.

| **Response  variable** | **Transformation** | **Fraction** | **Auto- correlation  structure** | **Explanatory  variable** | **Estimate** | **SE±** | **DF** | **t-value** | **p-value** |
| --- | --- | --- | --- | --- | --- | --- | --- | --- | --- |
| **APase** |  |  |  |  |  |  |  |  |  |
|  | log_10_ | <0.2 µm | – | Intercept | 0.478 | 0.076 | 74 | 6.237 | <0.0001 |
|  |  |  |  | **Temperature** | **0.031** | **0.004** | **74** | **6.610** | **<0.0001** |
|  |  |  |  | **Chlorophyll-a (µg l-1)** | **0.045** | **0.009** | **74** | **4.635** | **<0.0001** |
|  | sqrt | 0.2–3 µm | corAR1 | Intercept | –0.397 | 0.824 | 76 | –0.482 | 0.6307 |
|  |  |  |  | **DIP (µM)** | 3.043 | 0.479 | 76 | 6.348 | **<0.0001** |
|  |  |  |  | **POP (µM)** | 10.484 | 1.512 | 76 | 6.932 | **<0.0001** |
|  |  |  |  | Chlorophyll-a (µg l-1) | –0.390 | 0.123 | 76 | –3.152 | 0.0023 |
|  |  |  |  | Temperature | –0.089 | 0.043 | 76 | –2.051 | 0.0437 |
|  | log_10_ | >3 µm | – | Intercept | 0.498 | 0.526 | 77 | 0.947 | 0.3465 |
|  |  |  |  | **POP (µM)** | **2.498** | **0.485** | **77** | **5.141** | **<0.0001** |
|  |  |  |  | POC:POP | 0.005 | 0.001 | 77 | 3.090 | 0.0028 |
|  |  |  |  | Temperature | –0.047 | 0.018 | 77 | –2.595 | 0.0113 |
|  |  |  |  | DOP (µM) | –1.513 | 0.594 | 77 | –2.544 | 0.0129 |
| **LAPase** |  |  |  |  |  |  |  |  |  |
|  |  | <0.2 µm | corARMA | Intercept | 59.541 | 7.359 | 71 | 8.090 | <0.0001 |
|  |  |  |  | **NH4 µM** | **32.039** | **8.890** | **71** | **3.603** | **<0.0001** |
|  |  |  |  | **Temperature** | **–1.836** | **0.411** | **71** | **–4.468** | **<0.0001** |
|  | log_10_ | 0.2–3 µm | corARMA | Intercept | 0.000 | 0.000 | 76 | 0.833 | 0.4070 |
|  |  | (cell-specific) |  | Cell-specific BPT | 990.618 | 262.672 | 76 | 3.771 | 0.0003 |
|  |  |  |  | Temperature | 0.000 | 0.000 | 76 | 3.002 | 0.0036 |
|  |  |  |  | DOC (µM) | 0.000 | 0.000 | 76 | –2.093 | 0.0396 |
|  | sqrt | >3 µm | corAR1 | Intercept | 32.035 | 5.929 | 76 | 5.402 | <0.0001 |
|  |  |  |  | NH4 µM | –18.390 | 7.251 | 76 | –2.536 | 0.0133 |
|  |  |  |  | Temperature | –0.557 | 0.257 | 76 | –2.165 | 0.0335 |
|  |  |  |  | POC:PON ratio | –0.584 | 0.289 | 76 | –2.017 | 0.0472 |
| **AGase** |  |  |  |  |  |  |  |  |  |
|  |  | 0.2–3 µm | – | Intercept | 0.000 | 0.000 | 74 | 1.881 | 0.063 |
|  |  | (cell-specific) |  | Temperature | 0.000 | 0.000 | 74 | 4.022 | 0.0001 |
|  |  |  |  | DOC (µM) | 0.000 | 0.000 | 74 | –3.446 | 0.0009 |
|  |  |  |  | Chlorophyll-a (µg l-1) | 0.000 | 0.000 | 74 | –2.097 | 0.0048 |
|  |  |  |  | Cell-specific BPT | 11.534 | 4.243 | 74 | 2.718 | 0.0082 |
|  | sqrt | >3 µm | corAR1 | Intercept | 0.659 | 0.785 | 72 | 0.839 | 0.4041 |
|  |  |  |  | Temperature | –0.061 | 0.027 | 72 | –2.259 | 0.0269 |
|  |  |  |  | DOC (µM) | 0.003 | 0.001 | 72 | 2.192 | 0.0316 |
| **BGase** |  |  |  |  |  |  |  |  |  |
|  |  | 0.2–3 µm | – | Intercept | 0.000 | 0.000 | 75 | 0.752 | 0.453 |
|  |  | (cell-specific) |  | **Temperature** | **0.000** | **0.000** | **75** | **4.681** | **<0.0001** |
|  |  |  |  | Cell-specific BPT | 19.775 | 5.786 | 75 | 3.417 | 0.0010 |
|  |  |  |  | Chlorophyll-a (µg l-1) | 0.000 | 0.000 | 75 | –2.600 | 0.0110 |
|  |  |  |  | DOC:DON ratio | 0.000 | 0.000 | 75 | –2.871 | 0.0053 |

**Table S2.** List of significant results of linear mixed effects models investigating treatment effect. The table shows pairwise comparison between treatments conducted with the *emmeans* package in R.

| **Response  variable** | **Transformation** | **Fraction** | **Auto- correlation  structure** | **Contrast** | **Estimate** | **SE** | **DF** | **t-value** | **p-value** |
| --- | --- | --- | --- | --- | --- | --- | --- | --- | --- |
| **AGase** | sqrt | >3 µm | corAR1 | CTRL - NC | –0.616 | 0.168 | 8 | –3.674 | 0.0259 |
|  |  |  |  | C - NC | –0.780 | 0.166 | 8 | –4.697 | 0.0067 |
|  |  |  |  | N - NC | –0.632 | 0.166 | 8 | –3.808 | 0.0216 |
| **BGase** |  |  |  |  |  |  |  |  |  |
|  | log_10_ | >3 µm | corARMA | CTRL - NC | –0.374 | 0.094 | 8 | –3.953 | 0.0177 |
| **LAPase** |  |  |  |  |  |  |  |  |  |
|  |  | 0.2–3 µm | corARMA | N – NC | 2.5*10^-5^ | 7.2*10^-6^ | 8 | 3.434 | 0.0362 |
